# Ubiquitin modulates 26S proteasome conformational dynamics and promotes substrate degradation

**DOI:** 10.1101/2021.08.18.456915

**Authors:** Erik Jonsson, Zaw Min Htet, Jared A.M. Bard, Ken C. Dong, Andreas Martin

## Abstract

The 26S proteasome is the major ATP-dependent protease in eukaryotic cells, where it catalyzes the degradation of thousands of proteins for general homeostasis and the control of vital processes. It specifically recognizes appropriate substrates through attached ubiquitin chains and uses its ATPase motor for mechanical unfolding and translocation into a proteolytic chamber. Here, we used single-molecule Förster Resonance Energy Transfer (FRET) measurements to provide unprecedented insights into the mechanisms of selective substrate engagement, ATP-dependent degradation, and the regulation of these processes by ubiquitin chains. Our assays revealed the proteasome conformational dynamics and allowed monitoring individual substrates as they progress through the central channel during degradation. We found that rapid transitions between engagement- and processing-competent conformations of the proteasome control substrate access to the ATPase motor. Ubiquitin-chain binding functions as an allosteric regulator to slow these transitions, stabilize the engagement-competent state, and facilitate degradation initiation. The global conformational transitions cease upon substrate engagement, and except for apparent motor slips when encountering stably folded domains, the proteasome remains in processing-competent states for substrate translocation and unfolding, which is further accelerated by ubiquitin chains. Our studies revealed the dependence of ATP-dependent substrate degradation on the conformational dynamics of the proteasome and its allosteric regulation by ubiquitin chains, which ensure substrate selectivity and prioritization in a crowded cellular environment.

## INTRODUCTION

Eukaryotic cell viability critically depends on the 26S proteasome, which is responsible for protein homeostasis, quality control, and the degradation of numerous regulatory proteins ^1,2^. This major protease of the AAA+ (ATPases Associated with diverse cellular Activities) family recognizes protein substrates that have been modified with lysine-attached poly-ubiquitin chains, deubiquitinates them, and uses its ATPase motor for mechanical unfolding and translocation into an internal chamber for proteolytic cleavage ^3^. To degrade thousands of diverse proteins in a controlled manner, the 26S proteasome needs to combine rapid substrate processing and high promiscuity, with sufficient selectivity to avoid unregulated proteolysis of cellular proteins in general. This regulation is accomplished by the intricate architecture of the proteasome and its major conformational changes that help coordinate individual degradation steps ^4^, as well as the bipartite character of a substrate’s degradation signal, which consists of a poly-ubiquitin modification for targeting to the proteasome and an unstructured initiation region of at least 20-25 residues for engagement by the proteasomal ATPase motor ^4–10^.

Substrate cleavage by the proteasome occurs in the internal degradation chamber of the barrel-shaped 20S core peptidase, with access through axial pores being controlled by the 19S regulatory particle ^11–13^. This regulatory particle consists of the base and lid subcomplexes, and functions in ubiquitin-mediated substrate recognition, deubiquitination, ATP-dependent mechanical unfolding, and translocation into the 20S core. The base subcomplex contains 10 subunits, including three ubiquitin receptors, Rpn1 ^14^, Rpn10 ^15^, and Rpn13 ^16^, and a heterohexameric ATPase motor formed by six distinct subunits, Rpt1-Rpt6, with an N-terminal domain ring (N-ring) stacked on top of an AAA-domain ring ^14,17–19^. After ubiquitin binding to a receptor, the flexible initiation region of the substrate must reach through the N-ring and into the ATPase ring to engage with pore loops that transduce ATP hydrolysis into mechanical pulling for substrate unfolding and translocation into the 20S core ^4,20,21^. The 9-subunit lid subcomplex is bound to one side of the base and contains the deubiquitinating enzyme (DUB) Rpn11 ^22–26^, which catalyzes the co-translocational deubiquitination of substrates prior to their entry into the AAA+ motor ^27^.

Previous structural studies identified multiple proteasome conformations that differ in the relative orientation and interactions of the lid, base, and core subcomplexes ^28–36^ and provide a global framework for the mechanism of the 26S proteasome (Fig 1a). Dominant in the absence of substrate is the engagement-competent s1 state, in which the ATPase ring is not coaxially aligned with the core peptidase and Rpn11 is offset from the central processing channel, allowing substrate access to the entrance of the AAA+ motor. In the presence of substrate or the non-hydrolyzable ATP analog ATPγS, the proteasome adopts various processing states named s2-s6, or collectively non-s1 ^4,28,30,32,33,35,36^, in which the N-ring, ATPase ring, and core peptidase are coaxially aligned to form a continuous channel for efficient substrate translocation. Furthermore, Rpn11 is moved to a centrally aligned position that partially obstructs the motor entrance and thus facilitates *en bloc* ubiquitin-chain removal from substrates as they are translocated into the motor (Fig. 1a) ^27,35,36^. Structures of substrate-bound proteasomes in non-s1 states show four or five Rpt subunits with their pore loops contacting the substrate polypeptide and adopting different spiral-staircase arrangements around the hexameric ring ^35,36^, suggesting that translocation occurs by a hand-over-hand mechanism with a basic step size of two amino acids per hydrolyzed ATP, similar to related AAA+ protein translocases ^37–39^.

**Figure 1:**
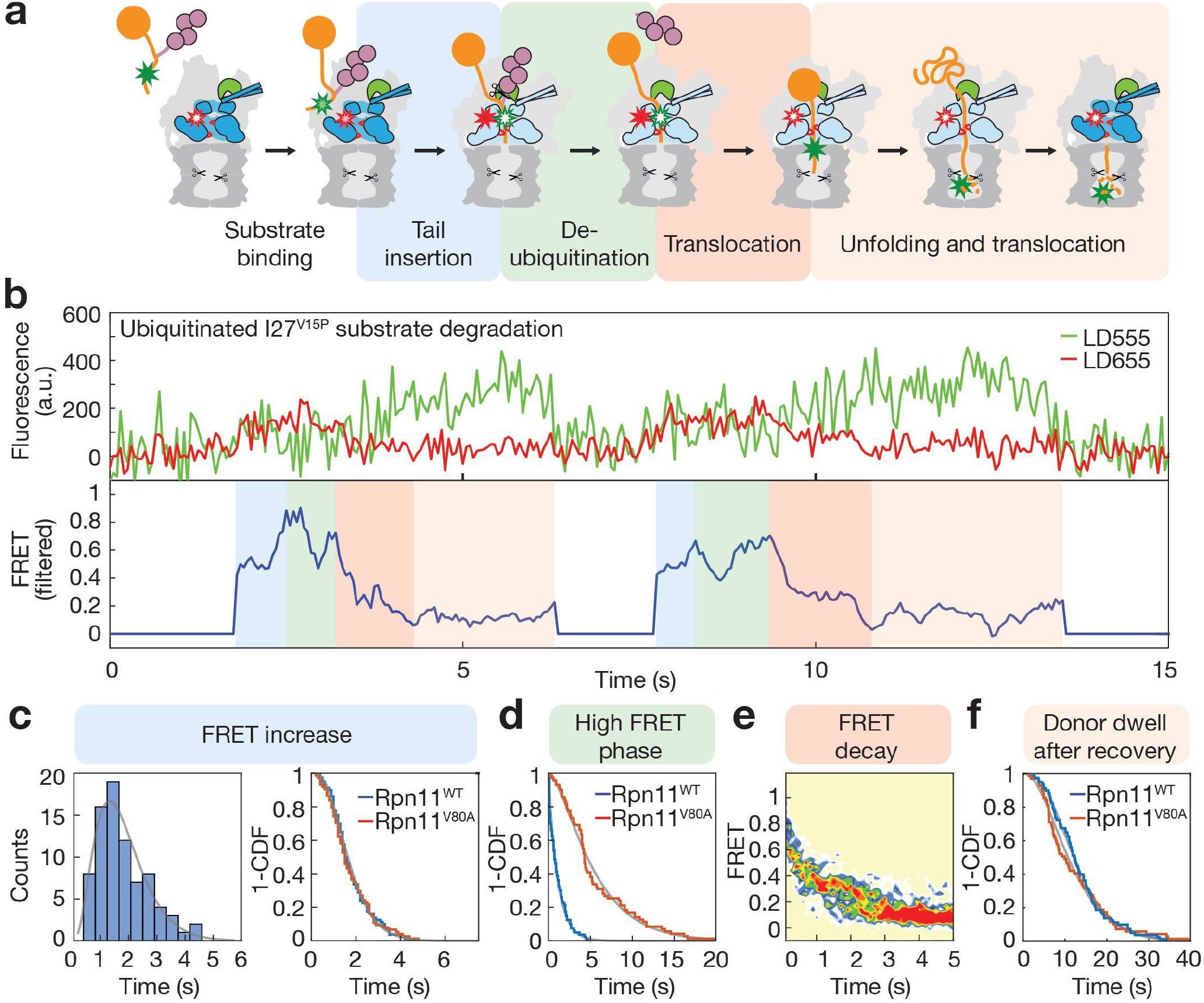
Direct observation of substrate processing. **a)** Cartoon representation depicting the sequential stages for the proteasomal degradation of an ubiquitinated substrate. The 20S core peptidase is shown in dark gray and the 19S regulatory particle in light gray, with Rpn11 highlighted in green, the Rpt1-6 ATPase in dark blue for the s1 conformational state and light blue for non-s1 states, and pore-1 loops shown as red hooks. The substrate is depicted in orange with the ubiquitin chain in purple. Green and red stars represent the substrate-attached donor and proteasome-attached acceptor dyes, respectively, with the filling indicating their expected fluorescence upon FRET during the different stages of substrate processing. **b**) Representative trace showing two consecutive events of ubiquitinated titin I27^V15P^ degradation by a single immobilized wild-type proteasome, with the donor fluorescence signal in green, acceptor fluorescence in red, and calculated filtered FRET in blue. The FRET and fluorescence signals show four phases that are shaded in different colors according to the substrate-processing steps shown in panel a). **c)** Left, histogram for the duration of the FRET-increase phase during degradation of ubiquitinated I27^V15P^ substrate, with the fit to a gamma distribution shown as a grey line (N = 80). Right, 1-CDF (cumulative distribution function) or survival plots for the FRET-increase phases during the degradation of ubiquitinated I27^V15P^ substrate by wild-type (blue) and Rpn11^V80A^-mutant (red) proteasomes (N = 80 and 75, respectively). Fit lines in grey are based on the cumulative distribution of the gamma function derived from fitting the dwell-time distributions. **d**) Survival plots for the high-FRET phase during degradation of ubiquitinated I27^V15P^ substrate by wild-type (blue) and Rpn11^V80A^-mutant (red) proteasome (N = 80 and 71, respectively). **e**) Contour plot for multiple FRET decay trajectories during degradation of ubiquitinated I27^V15P^ substrate by wild-type proteasome. Trajectories were synchronized using the point where the decay from the high-FRET value initiates (N = 53). **f**) Survival plot for the dwell times of donor dyes after fluorescence recovery during degradation of ubiquitinated I27^V15P^ substrate by wild-type (blue, N = 80) and Rpn11^V80A^-mutant proteasome (red, N = 69).

Previous ensemble-FRET-based measurements revealed that the major conformational changes from the s1 to non-s1 states are triggered by insertion and motor engagement of the substrate’s flexible initiation region (Fig. 1a) ^4^, and thus seems to play an important role in coordinating subsequent substrate-processing steps, including translocation-coupled deubiquitination. Interactions at the lid-base interface, in particular between the lid subunit Rpn5 and the Rpt3 ATPase, were found to stabilize the s1 state, and their disruption causes inhibited degradation due to compromised substrate engagement ^40^. Furthermore, the nucleotide state and hydrolysis activity of individual ATPase subunits, for instance Rpt4 and Rpt6, affect the proteasome’s conformational equilibrium ^40,41^.

Despite these previous insights, the detailed mechanisms underlying substrate engagement, deubiquitination, mechanical unfolding, and translocation, as well as the role of proteasome conformational transitions for the individual processing steps remain largely elusive. Furthermore, ubiquitin modifications were observed to facilitate substrate degradation ^42,43^, yet their effects beyond substrate recruitment, for instance on degradation initiation, mechanical processing, or the proteasome conformational states are unknown.

Here, we used single-molecule FRET measurements to monitor the conformational dynamics of the proteasome and observe individual substrate molecules on their way through the regulatory particle toward degradation. These experiments gave new insights into the velocity of translocation and translocation-coupled deubiquitination, and revealed the rapid conformational switching of the proteasome regulatory particle that depends on a network of interactions at the lid-base interface and plays an important role in selective substrate engagement. Furthermore, we uncovered how ubiquitin-chain binding to the proteasome allosterically affects these conformational transitions and facilitates degradation initiation as well as mechanical unfolding, providing exciting new insights into the mechanism and regulation of ubiquitin-dependent substrate degradation by the proteasome.

## RESULTS

### Substrate progression through the 26S proteasome

We used single-molecule FRET to observe individual substrates as they translocate through the proteasome regulatory particle on their way to degradation. Our model substrates consisted of a titin-I27 domain, either wild type or carrying the destabilizing mutations V13P and V15P ^4,44^, and a C-terminal cyclin-B-derived unstructured initiation region that was labeled with a LD555- or Cy3-donor dye and ubiquitinated on a single lysine residue (Supp. Fig 1b). Reconstituted proteasomes with a LD655-acceptor-dye labeled azido-phenylalanine in the linker between the N-domain and the AAA+ domain of the Rpt1 ATPase subunit were immobilized to the surface of a microscope cover slip (Supp. Fig. 1a), substrates were added to the reaction chamber, and fluorescence signals were detected in a total internal reflection fluorescence (TIRF) microscope (Supp. Fig. 2). By monitoring variations in fluorescence intensity as the distance between donor and acceptor dyes changes, we could observe individual substrate molecules as they enter the proteasome and progress through the central channel during the various stages of processing (Fig. 1a). Proteasomes often showed several sequential degradation events, and a representative trace with two consecutive degradations of ubiquitinated titin I27^V15P^ by a single proteasome is depicted in Fig. 1b. All degradation events exhibit an evolution of the FRET signal, starting from an intermediate level, followed by an increase and a short dwell in a high FRET state, before gradually decaying to a background level. The FRET decay occurs with a concomitant increase in donor fluorescence that persists for a while before disappearing in a single step, which confirms the observation of a single labeled substrate molecule. These FRET and fluorescence profiles are consistent with a substrate-attached donor dye localizing near the regulatory particle upon substrate binding, entering the central processing channel of the ATPase motor, translocating towards, past, and away from the motor-attached acceptor dye, and then progressing into the 20S core particle, where substrate cleavage and diffusion of the donor-labeled peptide out of the proteolytic chamber leads to disappearance of the donor signal (Fig. 1a,b). Measuring the signal increase from the start of the reaction to the point of highest FRET gave a time constant of τ_ins_ = 1.8 ± 0.1 s (N = 80; Fig. 1c), which is in excellent agreement with previous bulk measurements of substrate tail insertion ^4^ and indicates full activity of the surface-immobilized proteasomes. This tail insertion includes the passive diffusion of the substrate’s C-terminal initiation region into the central channel, the engagement by the ATPase motor, and the onset of translocation. Based on our structure of the substrate-engaged proteasome ^35^, we placed the donor dye on the substrate’s unstructured tail such that it localizes between the N-ring and ATPase ring, causing a high-FRET signal, when the tail is fully inserted and the substrate-linked ubiquitin chain reaches the Rpn11 deubiquitinase above the N-ring (Supp. Fig. 1b). Deubiquitination is therefore expected to occur during the high-FRET dwell phase, which we measured to last only for τ_DUB_ = 1.1 ± 0.2 s (N = 80; Fig. 1d, Supp. Fig. 3a). We can conclude that degradation-coupled ubiquitin-chain removal proceeds at least 4 times faster than the 4.6 s we previously determined in bulk measurements with a FRET-based deubiquitination assay that did not allow a separation from tail insertion or translocation immediately after deubiquitination ^4^. To verify this assignment of deubiquitination during the high-FRET phase, we introduced the Rpn11 V80A point mutation that was previously shown in bulk measurements to slow down substrate deubiquitination by ~ 4-fold ^27^. Consistently, we observed extended high-FRET dwells of τ_DUB_ = 5.5 ± 0.5 s (N = 71; Fig. 1d, Supp. Fig.3b), whereas other phases of substrate processing, such as tail insertion, remained unaffected (Fig. 1c, right).

After ubiquitin-chain removal, the proteasomal ATPase motor is expected to freely translocate the 24 residues between the deubiquitinated lysine and the titin I27 domain in our model substrate (Supp. Fig. 1b), leading to a gradual decay and loss of FRET before the folded domain reaches the narrow entrance to the N-ring (Fig. 1e). Based on the structure of the substrate-bound proteasome ^35^ and the geometry of our labeled constructs, the observed FRET decay would be consistent with an advancement of the donor dye by ~ 62 Å in the central channel and a corresponding translocation of ~ 22-24 residues. That this FRET decay reflects unobstructed substrate translocation prior to titin unfolding is supported by the observation that titin I27^V15P^ and the more strongly destabilized I27^V13P/V15P^ double-mutant substrate show similar kinetics (Supp. Fig. 3c). Fitting the FRET- decay during I27^V15P^ degradation to a single exponential revealed a time constant of 0.78 s ± 0.1 s (Supp. Fig. 3d), corresponding to a translocation rate of *k_trans_* ~ 30 aa s^−1^. The proteasome thus appears to thread a polypeptide substrate an order of magnitude faster than the ~ 2 - 3 aa s^−1^ suggested by a structure-derived step size of 2 aa ATP^−1 35^ and a bulk ATP-hydrolysis activity of 1.2 ATP s^−1 40^. Either bulk ATPase measurements do not accurately reflect the proteasome’s hydrolysis rate during unobstructed substrate translocation or AAA+ motors take larger steps than implied by their static substrate-bound structures ^35,37–39^. In fact, much larger steps of 1 - 4 nm, equivalent to 5 – 20 aa, were previously observed in single-molecule optical tweezers experiments with the related bacterial protease ClpXP ^45–48^.

By the time the motor attempts to mechanically unfold the titin domain, the donor dye has proceeded through the ATPase ring to a range within the 20S core particle that is no longer amenable to efficient energy transfer. Nevertheless, the prolonged persistence of the donor fluorescence at the position of the proteasome in our TIRF measurements reports on the approximate time required for substrate unfolding, translocation to the peptidase active sites, cleavage, and release of the peptides from the proteolytic chamber, and we detected a donor dwell of 13.3 ± 0.8 s for this complete processing of the I27^V15P^ substrate (Fig. 1f). Rpn11^V80A^-mutant proteasomes showed a similar donor dwell (Fig. 1f), confirming that the Rpn11 activity has no effect on the processing steps after deubiquitination.

### Observing the proteasome conformational dynamics

Numerous cryo-EM studies indicated that upon substrate engagement by the AAA+ motor, the proteasome transitions from an engagement-competent s1 state with an accessible motor entrance to non-s1 or processing-competent states, in which Rpn11 is positioned right above the central channel for efficient co-translational deubiquitination ^28,32,33,35,36^. During this global conformational switch, the distance between the lid subunit Rpn9 and the N-terminal coiled-coil of the Rpt4/Rpt5 ATPase pair decreases by ~ 30 Å. We previously developed a FRET assay with dyes attached in those positions to monitor the proteasome conformational change upon substrate engagement in bulk measurements ^4^. Here, to investigate the conformational dynamics of individual proteasomes, we optimized this FRET system for single-molecule experiments with a LD555 donor dye on Rpn9 and a LD655 acceptor dye on Rpt5 (Fig. 2a, Supp. Fig. 1a). A representative trace for the substrate-free proteasome in the presence of ATP is depicted in figure 2b, indicating dynamic switching between two conformational states (Supp. Fig. 4a). Based on the positioning of the dyes, we assign the low-FRET state to s1 and the high-FRET state to non-s1 conformations. Previous structural studies revealed that the individual non-s1 states differ in their spiral-staircase arrangements of the six ATPase subunits, but exhibit no global changes in the relative orientation of the lid and base subcomplexes ^32,35,36^. We therefore did not expect the FRET probes to distinguish between individual non-s1 states, and indeed our measurements are consistent with a two-state system (Supp. Fig. 4b). The FRET histogram shows that the low-FRET s1 state is predominant, whereas the high-FRET non-s1 states are only transiently visited under these conditions (Fig. 2c). The dwell times in each state could be determined through Hidden Markov modeling and allowed us to derive the kinetics for conformational switching, with k_s1_ = 1.2 ± 0.1 s^−1^ and k_non-s1_ = 4.5 ± 0.1 s^−1^ for the s1-to-non-s1 and non-s1-to-s1 transitions, respectively (Supp. Fig. 4c, d). Addition of the non-hydrolyzable ATP analog, ATPγS, shifts the histogram completely to the high-FRET non-s1 states (Fig. 2c, Supp. Fig. 6a), which is in agreement with previous cryo-EM studies ^29^ and bulk measurements ^4^, and confirms our conformational assignment of FRET levels.

**Figure 2:**
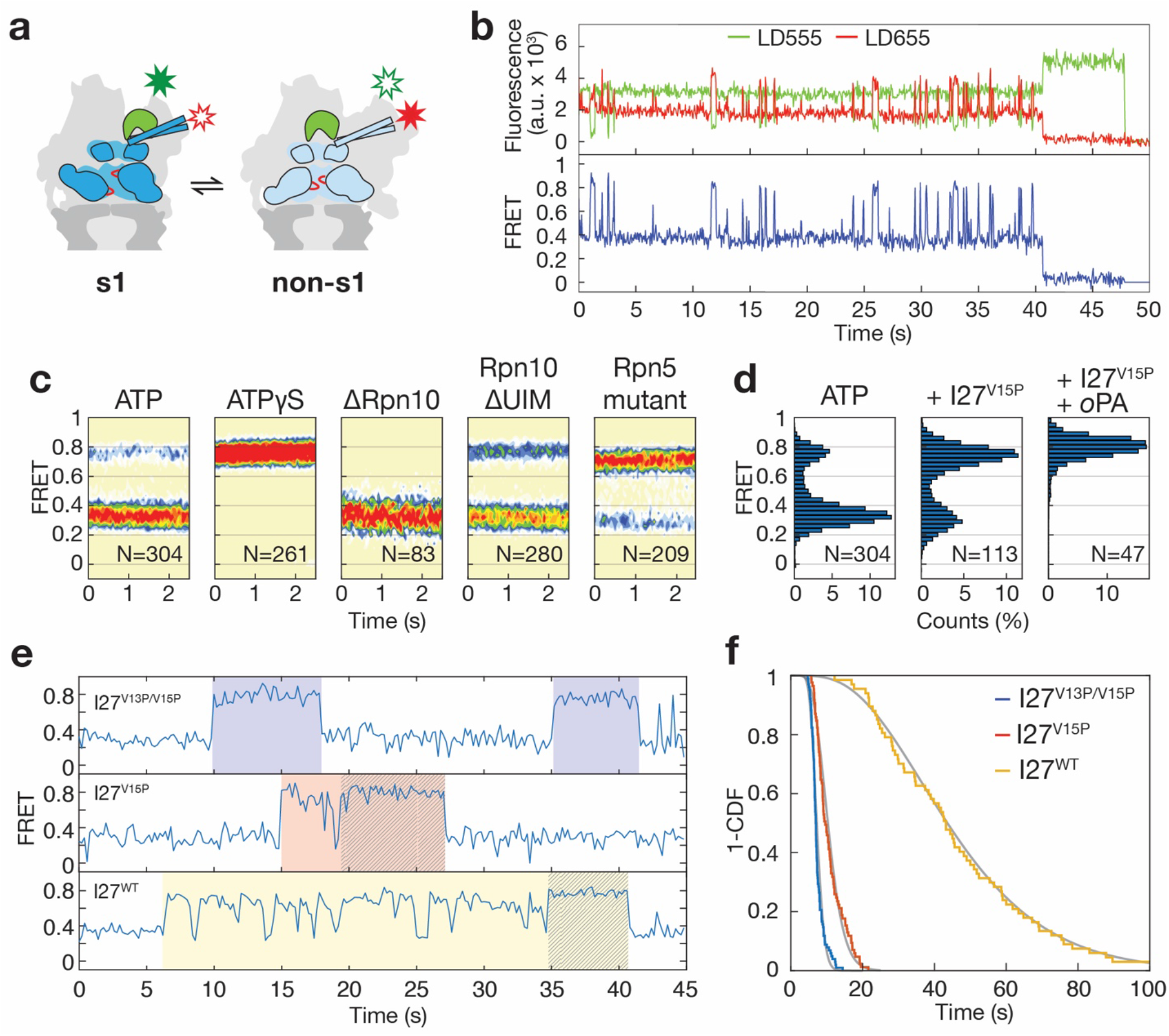
Conformational dynamics of the proteasome. **a**) Cartoon representation for the conformational transition between the engagement-competent s1 and the processing-competent non-s1 states. A lid-attached donor dye (green star) and base-attached acceptor dye (red star) allow monitoring these transitions by FRET in a conformational-change assay (low FRET in s1, high FRET in non-s1). **b**) Representative trace for a wild-type proteasome in ATP exhibits dynamic switching between low-FRET and high-FRET states, with raw donor and acceptor fluorescence signals shown in green and red, respectively, and the calculated FRET value in blue. **c**) Contour plots generated from time-binned histograms of multiple traces indicate the relative occupancies of low-FRET and high-FRET states for the 26S proteasome in the presence of ATP and the non-hydrolyzable ATPγS, upon deletion of the entire Rpn10 or its UIM, and upon insertion of s1-state destabilizing mutations in Rpn5. **d**) FRET distribution of wild-type proteasome in the presence of ATP (left), upon addition of 400 nM ubiquitinated titin I27^V15P^ substrate (middle), and addition of substrate and the Rpn11 inhibitor 1,10-phenanthroline (*o*PA) at 3 mM (right). **e**) Representative FRET traces for the conformational dynamics of the proteasome during SspB-mediated degradation of the I27^V13P/V15P^, I27^V15P^, and I27^WT^ substrates with different thermodynamic stabilities (traces for I27^V13P/V15P^ and I27^V15P^ were down-sampled to 5 Hz to match I27^WT^). Substrate-processing dwells are highlighted by shading in blue, red, and yellow, and their length as well as the frequency of excursion to the low-FRET s1 state increase with increasing substrate stability. The last 3-5 seconds of traces show an excursion-free high-FRET state that is indicated by hatching. **f**) Survival plots for the substrate-processing dwells during SspB-mediated degradation of the I27^V13P/V15P^ (N = 102), I27^V15P^ (N = 85), and I27^WT^ (N = 67) substrate variants. Fit lines in grey are based on the cumulative distribution of the gamma function derived from fitting the dwell-time distributions.

Low ATP concentrations bias the conformational equilibrium towards non-s1 states, and the absence of nucleotide leads to an almost complete shift (Supp. Fig. 5a, b), similar to the scenario observed in the presence of the non-hydrolyzable ATPγS. This finding suggests that establishing all Rpt subunits in identical nucleotide states, i.e. empty or ATPγS-bound, favors non-s1 conformations, whereas the presence of various nucleotide states in the Rpt hexamer during active ATP hydrolysis may provide strains that stabilize the more distorted s1 state, at least in the absence of substrate. The presence of only ADP, however, destabilizes the proteasome holoenzyme and leads to the dissociation of the regulatory particle from the 20S core, as indicated by the lack of co-localized lid and base subcomplexes in our conformational change assay (Supp. Fig. 5c).

We also analyzed proteasomes containing mutations in the VTENKIF motif of the lid subunit Rpn5 that forms s1-state-specific contacts with the ATPase subunit Rpt3. Disruption of these contacts was previously found to inhibit substrate degradation by interfering with initiation ^40^. We observed that in the presence of ATP, the Rpn5-mutant proteasomes not only exhibit a bias away from the s1 state (Fig. 2c), but also switch much more rapidly between conformations (k_s1_ = 2.5 ± 0.1 s^−1^ and k_non-s1_ = 4.2 ± 0.1 s^−1^, Supp. Fig. 6d, Table 2). These results are consistent with a mechanism of degradation inhibition by which accelerated conformational transitions and/or a shift in the equilibrium toward non-s1 states with an Rpn11-obstructed motor entrance hinder substrate engagement ^40^. This behavior of the Rpn5 mutant highlights the importance of the engagement-competent s1 state being long-lived enough to allow substrate insertion into the central channel, and it emphasizes the critical role of lid-base interactions in controlling the dynamics of the proteasome during various stages of degradation.

Interestingly, holoenzymes reconstituted in the absence of the ubiquitin receptor Rpn10, which bridges the lid and base subcomplexes, show the conformational equilibrium shifted towards a low-FRET state that according to previous EM studies likely reflects the s1 conformation (Fig. 2c, Supp. Fig. 6b) ^49^. Consistent with these effects in ATP, ΔRpn10 proteasomes with bound ATPγS still show a bimodal distribution of the FRET histogram and a significant population of s1-state particles (Supp. Fig. 7), suggesting that Rpn10 plays a role in stabilizing non-s1 states, at least in the absence of substrate. This non-s1 stabilization appears to be contributed by Rpn10’s globular VWA domain, as deletion of Rpn10’s C-terminal ubiquitin-interacting motif (UIM) alone shifts the equilibrium compared to wild-type proteasomes toward non-s1 states (Fig. 2c, Supp. Fig. 6c). Together, our data indicate the presence of an intricate system of interactions at the lid-base interface that differentially stabilize the s1 and non-1 states, and thus allows for dynamic conformational transitions and equilibrium shifts depending on the stage of substrate processing.

### Conformational response to substrate engagement and processing

The addition of ubiquitinated substrate at saturating concentrations shifted the proteasome conformational equilibrium toward non-s1 states (Fig. 2d), in agreement with previous findings that substrate engagement induces a s1-to-non-s1 transition ^4,28^. The observed minor s1 peak in the histogram is explained by the asynchronicity of individual particles and the fact that proteasomes appear to “idle” in the s1 state between degradation events. Consistently, the conformational equilibrium could be fully shifted to high-FRET states by stalling substrate degradation at the deubiquitination phase through addition of the Rpn11 inhibitor 1,10-phenanthroline (*o*PA) (Fig. 2d) ^27^. To distinguish between potential effects of ubiquitin binding and substrate degradation on proteasome conformations, we used a previously developed ubiquitin-independent substrate delivery system. In this system, the bacterial adaptor SspB is fused to the N-terminus of the Rpt2 ATPase subunit, which allows the recruitment and degradation of non-ubiquitinated substrates that contain the SspB-interacting ssrA tag in their unstructured initiation region (Supp. Fig. 8a, b) ^50^. FRET traces in the conformational change assay revealed that proteasomes with SspB-delivered or ubiquitinated substrates exhibit similar prolonged dwells in the high FRET state that likely represent individual substrate-processing events (Fig. 2d, Supp. Fig. 8c-e). This interpretation is supported by the observed correlation between the length of these high-FRET dwells and the thermodynamic stability of the substrate’s I27 domain. SspB-delivered wild-type I27 substrate showed significantly longer processing dwells (τ_deg_ = 47 ± 4 s) than the I27^V15P^ single-mutant (τ_deg_ = 10.7 ± 0.7 s) or the least stable I27^V13P/V15P^ double-mutant substrate (τ_deg_ = 7.3 ± 0.2 s, Fig. 2e, f, Supp. Fig. 9 and Table 3). Given that the conformational switch to high-FRET non-s1 states is induced by substrate engagement, it can be assumed that the proteasome stably switches back to the s1 state after translocation has been completed and the substrate terminus has cleared the ATPase ring. The high-FRET dwells in the conformational change assay therefore represent an accurate readout for the total time of substrate processing after engagement, and the derived time constants for all three substrate variants are indeed in excellent agreement with previous anisotropy-based measurements of single-turnover degradation in bulk ^4^. Based on a velocity of 30 aa s^−1^ for unobstructed translocation, it would take ~ 4.6 s to completely thread the ~ 138 residues that reside above or within the ATPase ring after initiation of our model substrates (Supp. Fig. 1b). By subtracting this translocation time from the total time of substrate processing, we can estimate the unfolding time constants as τ_unfold_ = 42 s for wild-type I27, τ_unfold_ = 6.1 s for I27^V15P^, and τ_unfold_ = 2.7 s for I27^V13P/V15P^.

Interestingly, the high-FRET dwells during substrate processing show intermittent, brief returns to the low-FRET s1 state, with a frequency that depends on the thermodynamic stability of the substrate (Fig. 2e, Supp. Fig. 9). While the strongly destabilized I27^V13P/V15P^ variant shows barely any of these transitions to the s1 state, they are more frequent (0.7 ± 0.2 per processing) for I27^V15P^ and very prominent (11 ± 2 per processing) during the motor’s attempts to unfold wild-type I27. However, for all substrate variants, the s1-returns are noticeably absent from the last 3 - 5 seconds of processing (Fig. 2e, Supp. Fig. 9b,c), which likely reflect the unobstructed translocation of the substrate polypeptide after successful unfolding and is consistent with the translocation rate we determined above. Intermittent s1-returns during the unfolding phase may thus represent slippage events, where the pore loops struggle with a high unfolding-energy barrier and lose grip of the polypeptide, leading to a transient elimination of the substrate-induced stabilization of non-s1 states. Such slippage would be consistent with our previous findings that ATP hydrolysis, and presumably motor movements, do not cease when substrate translocation stalls at a barrier ^12^. Importantly, the uninterrupted high-FRET dwells during processing of I27^V13P/V15P^ or translocation of the unfolded polypeptides contradict a previously proposed ratcheting model, in which substrate translocation is driven by obligatory, repetitive transitions between s1 and non-s1 states. Instead, the proteasome appears to switch out of the s1 state upon substrate engagement and then cycle through the various non-s1 states with different spiral-staircase arrangements of ATPase subunits to propel substrates by some kind of hand-over-hand mechanism, not requiring any returns to s1 ^35,36^. We also observed that ΔRpn10 and Rpn5^VTENKIF^-mutant proteasomes exhibit high-FRET dwells during I27^V13P/V15P^ substrate degradation that are of similar persistence and length (τ_deg_ = 7.9 ± 0.3 s and τ_deg_ = 8.4 ± 0.3 s, respectively) as those of wild-type proteasomes (Supp. Fig. 8c,d). Although these mutations shift the conformational equilibrium in the absence of substrate, they do not affect processing once a substrate is engaged, which is consistent with our model that Rpt pore-loop interactions with the substrate polypeptide dominate the conformational equilibrium over lid-base contacts or the effects of ATP binding and hydrolysis by the Rpt motor.

### Ubiquitin slows conformational switching and accelerates degradation

A major advantage of our SspB-mediated substrate-delivery system is that it allows us to directly measure potential allosteric effects of ubiquitin-chain binding to the proteasome on substrate degradation and the conformational dynamics. In bulk measurements, we observed a ~ 1.4-fold acceleration of titin-substrate degradation by unanchored K48-linked ubiquitin tetramers (K48-Ub_4_, Fig. 3a), which is consistent with a previous study reporting that substrate-attached ubiquitin chains affect proteasomal degradation ^42^. To determine which steps of degradation were modulated by ubiquitin-chain binding to the proteasome, we first analyzed the effects of unanchored K48-Ub_4_ in our FRET-based conformational dynamics assay. In the absence of substrate, addition of K48-Ub_4_ biased the proteasome conformational equilibrium towards the s1 state (Fig. 3b, Supp. Fig. 10, 11a). Linear and K63-linked ubiquitin chains showed similar effects (data not shown), suggesting that diverse ubiquitin-linkage types induce this common conformational response. Ubiquitin chains decrease the rate of the s1–to–non-s1 transition by ~ 3-fold, from 1.2 to 0.4 s^−1^, while having no significant effects on the return from non-s1 to the s1 state (Supp. Table 2). Ubiquitin binding thus stabilizes the proteasome in the engagement-competent s1 conformation with a well-accessible central channel, and we hypothesized that it promotes substrate insertion and degradation initiation.

**Figure 3:**
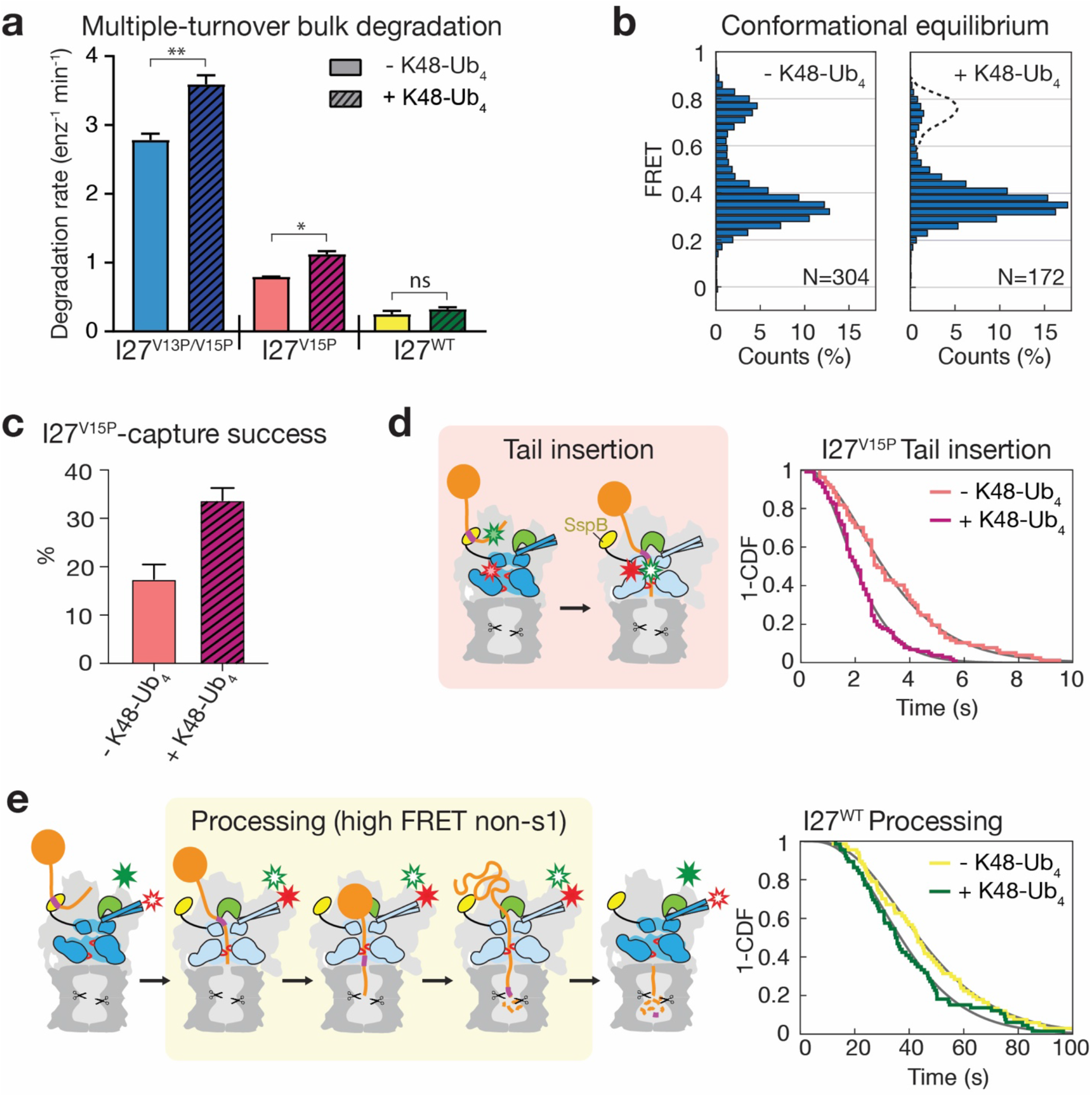
Ubiquitin chains modulate conformational dynamics and activate the proteasome. **a**) Rates for the multiple-turnover degradation of the I27^V13P/V15P^, I27^V15P^, and I27^WT^ substrates after SspB-mediated delivery to the proteasome in the absence (solid bars) and presence (hatched bars) of 10 μM unanchored K48-linked tetra-ubiquitin chains (K48-Ub_4_) (N = 3 technical replicates). Statistical significance was calculated using an unpaired two-tailed Welch’s t-test: **p = 0.0091; *p = 0.0185; ns, p = 0.2766. **b**) Histograms for the FRET-state distributions of wild-type proteasomes in the absence (left) and presence (right) of K48-Ub_4_. The dashed line on the right shows the distribution in the absence of K48-Ub_4_ for comparison. **c**) Substrate-capture success of the proteasome, as determined by the fractional rate of productive substrate-binding and degradation events relative to all substrate encounters for SspB-delivered I27^V15P^, in the absence (red) and presence (magenta) of K48-Ub_4_. **d**) Left, schematic for the tail insertion of SspB-delivered substrate. Right, survival plots for the tail-insertion times of SspB-delivered I27^V15P^ substrate in the absence (red, N = 77) and presence of K48-Ub_4_ (magenta, N = 102). Fit lines in grey are based on the cumulative distribution of the gamma function derived from fitting the dwell-time distributions. **e**) Times for the substrate-processing dwells during degradation of SspB-delivered I27^WT^ (yellow highlight in the schematic, left) are shown as survival plots (right) for the absence (yellow, N = 67) and presence of K48-Ub_4_ (green, N = 66).

To test this, we utilized our FRET-based substrate-processing assay to monitor SspB-mediated I27^V15P^ degradation and analyze the engagement efficiency or “capture success rate”, which is reflected by the number of complete processing events per total number of substrate encounters (example trace shown in Supp. Fig. 8b). The presence of ubiquitin chains nearly doubled this capture success, from 17 ± 3 % to 34 ± 3 % (Fig. 3c), a value that is very similar to the capture success for the ubiquitinated substrate (33 ± 2 %). Furthermore, we observed that proteasome-bound ubiquitin chains accelerate substrate-tail insertion, lowering the time constant from 3.4 s to 2.2 s, which approaches the value for ubiquitinated substrates (Fig. 3d, Supp. Table 1). This confirmed our hypothesis that the ubiquitin-mediated increase in lifetime of the s1 state to 2.5 s (k_s1_ = 0.4 s^−1^), and thus into the range of substrate tail-insertion kinetics, facilitates degradation initiation and increases capture success. The SspB-fused proteasomes used for these substrate-processing assays show a s1-to-non-s1 transition rate in the absence of ubiquitin chains that is ~ 30 % lower than for wild-type proteasomes (k_s1_ = 0.8 s^−1^ versus 1.2 s^−1^, Supp. Table 2), possibly due to steric effects, yet an identical rate of 0.4 s^−1^ in the presence of ubiquitin. The s1-state lifetime of SspB-fused proteasomes is therefore only 2-fold extended by ubiquitin-chain binding, whereas wild-type proteasomes show a 3-fold effect, which is expected to cause an even stronger ubiquitin-mediated acceleration of substrate-tail insertion than observed here with the SspB fusion. Substrate capture by the proteasome depends on the competition between the engagement of the polypeptide by the ATPase motor and the dissociation of a substrate-attached ubiquitin chain from proteasomal receptors or, for our ubiquitin-independent delivery system, the dissociation of the ssrA peptide motif from the proteasome-fused SspB adaptor. We measured the time constant for the dissociation of a tailless, ubiquitinated substrates from proteasomal receptors as τ_Ub-off_ = 0.61 ± 0.12 s (Supp. Fig. 11b), and a time constant of τ_ssrA-off_ = 0.32 s was previously determined for the dissociation of ssrA from SspB ^51^. Both dissociations occur much faster than the complete tail insertion and motor engagement of an ubiquitinated substrate (τ_ins_ = 1.8 s) or a SspB-delivered substrate in the presence of unanchored ubiquitin chains (τ_ins_ = 2.2 s). We therefore propose that a substrate’s flexible initiation region rapidly forms stabilizing interactions with the upper part of the processing channel that can prevent immediate substrate dissociation from the proteasome. This model is also consistent with our previous findings that the presence of a flexible initiation region increases the apparent substrate affinity for the proteasome by ~ 10-fold ^4^. The initial interactions with the channel are likely not stable enough to withstand a collision of the substrate polypeptide with Rpn11 when the proteasome transitions from the s1 to non-1 states, in which Rpn11 obstructs the central pore and leaves only a small gap above the N-ring for substrates to be translocated through (Fig. 1a, 2a). Successful substrate capture thus likely depends on a stable engagement with the pore loops of the ATPase motor before the proteasome switches to non-s1 states, which would explain why a ubiquitin-induced longer time constant for conformational switching facilitates substrate tail insertion and engagement, and causes an increase in the capture success rate.

To assess how ubiquitin chains modulate substrate unfolding and translocation, we again used the FRET-based conformational change assay with SspB-fused proteasomes. In the presence of K48-Ub_4_ chains, we observed high-FRET dwells of τ_deg_ = 40 ± 4 s for wild-type I27, τ_deg_ = 9.0 ± 0.3 s for I27^V15P^, and τ_deg_ = 7.0 ± 0.1 s for I27^V13P/V15P^, which is 7 s, 1.7 s, and 0.3 s, respectively, faster than in the absence of ubiquitin (Fig. 3e, Supp. Table 3). In the FRET-based substrate-tail insertion and processing assay, the FRET decays after degradation initiation indicate no significant differences in translocation velocity with and without K48-Ub_4_ (Supp. Fig. 11c), indicating that the observed acceleration of substrate processing is due to faster unfolding in the presence of ubiquitin chains. This is further supported by the observation that the extent of degradation acceleration correlates with the substrate’s thermodynamic stability. Although ubiquitin binding modulates the conformational dynamics of the proteasome in the absence of substrate, there were no discernable differences in the proteasome dynamics during substrate processing with and without ubiquitin chains (Supp. Fig. 12). S1 excursions, and thus potential slipping events of the motor, were equally frequent prior to successful unfolding, and the mechanism for more efficient mechanical substrate unraveling in the presence of ubiquitin chains remains unclear.

## CONCLUSION

Our single-molecule FRET-based assays revealed that the proteasome’s dynamic switching between the engagement-competent s1 state and the processing-competent non-s1 states controls access to the central channel and represents a selectivity filter for appropriate substrates that can rapidly enter and engage with the ATPase motor. Ubiquitin-chain binding to the proteasome functions as an allosteric regulator that slows down this conformational switching, stabilizes the s1 state, facilitates insertion of the substrate’s flexible initiation region, and consequently doubles the success rate of capturing a substrate for degradation. Although the presence of unanchored ubiquitin chains appeared to only moderately increase the rates for SspB-mediated substrate degradation under saturating conditions in bulk measurements, ubiquitin’s allosteric effects on degradation are expected to be larger for covalently attached chains and provide a significant advantage especially in the cellular environment, where ubiquitinated substrates have to compete with non-ubiquitinated proteins that may be present at local high concentrations near the proteasome due to molecular crowding. Upon substrate engagement, the proteasome switches to the non-s1 states for substrate unfolding and translocation, which occurs at a velocity of ~ 30 aa s^−1^ and thus about an order of magnitude faster than previously assumed based on the small translocation step size derived from substrate-bound structures. Co-translocational ubiquitin-chain removal by Rpn11 also happens very rapidly, at a rate ~ 50-fold higher than the maximal rate for translocation-independent ubiquitin cleavage ^27^. Deubiquitination therefore does not contribute to the overall degradation time, even for substrates with multiple ubiquitin modifications ^4^. The time for mechanical unfolding correlates with the substrate’s thermodynamic stability and is shortened by ubiquitin-chain binding to the proteasome. How ubiquitin chains accelerate substrate unfolding remains unclear and will have to be investigated in future studies, but contrary to a recently proposed model ^52^, the proteasome’s conformational dynamics or the relative stabilities of the engagement-competent s1 versus processing-competent non-s1 states do not seem to play a role. Our single-molecule FRET measurements provide unprecedented detail about ubiquitin-dependent substrate processing by the proteasome and laid the foundation for further studies of how AAA+ motor engage, translocate, and unfold their substrates, and how these processes are allosterically regulated by effectors like ubiquitin chains.

## Acknowledgements

We thank the members of the Martin lab for helpful discussions and Nico Stuurman for help with the microscope setup. This research was funded by the Howard Hughes Medical Institute (E.J., Z.M.H, K.C.D., and A.M.) and by the US National Institutes of Health (R01-GM094497 to A.M.). J.A.M.B. acknowledges support from NSF Graduate Research Fellowships.

## Author contributions

E.J. and A.M. designed experiments, E.J., Z.M.H., and J.A.M.B. performed single-molecule and biochemical measurements, and K.C.D. assisted with the preparation of materials. All authors contributed to the manuscript preparation.

## METHODS

### Purification and labeling of the lid and base subcomplexes

Recombinant proteasomes were purified, fluorescently labeled, and reconstituted as described previously ^4,53^. Three separate purifications (base, lid and core) are required to reconstitute 26S recombinant proteasomes. For recombinant expression of the yeast base subcomplex, *E. coli* BL21-star(DE3) (Invitrogen) was co-transformed with the plasmids pAM81, pAM82, and pAM83 (derived from pETDuet, pCOLADuet, and pACYCDuet, respectively), which code for Rpt1 - Rpt6 (including an amber codon TAG at the desired position for site-specific unnatural amino acid incorporation), Rpn1, Rpn2, Rpn13, the four base-assembly chaperones Rpn14, Hsm3, Nas2, and Nas6, and rare tRNAs. An additional fourth plasmid, pUltra_pAzFRS.2.t1_UAG-tRNA ^4^, coded the azido-phenylalanine (AzF) t-RNA synthase/t-RNA to allow unnatural amino acid incorporation. Cells were grown at 37 °C in 6 liters of terrific broth (Novagen) to an OD_600_ between 0.6 and 0.9, then pelleted and resuspended in 1 liter of terrific broth containing 2 mM AzF (Amatek Chemical), incubated at 30 °C for 30 minutes, induced with 1 mM IPTG for 5 hours at 30 °C, and then further incubated overnight at 16 °C. Cells were pelleted and resuspended in lysis buffer (60 mM HEPES, pH 7.6, 100 mM NaCl, 100 mM KCl, 10 mM MgCl2, 5% glycerol, and 20 mM imidazole) supplemented 1 mM ATP, lysozyme, benzonase (Novagen) and protease inhibitors (aprotinin, leupeptin, pepstatin and PMSF). The cells were lysed by sonication, the lysate was clarified by centrifugation for 30 minutes at 30,000 g, and the base subcomplex was fluorescently labeled and purified by a three-step procedure, with 0.5 mM ATP present in all buffers. First, the His-Rpt3 containing complexes were isolated using a 5 mL HisTrap FF crude column (Cytiva), and then fully assembled base complexes containing Flag-Rpt1 were selected for using anti-Flag M2 affinity resin (Sigma). After concentration (Amicon Ultracel-100K), the base subcomplexes were incubated with 150 μM 5,5’-dithiobis-(2-nitrobenzoic acid) (DTNB) for 10 min at room temperature to reversibly block surface exposed cysteines, before the AzF was reacted with 300 μM dibenzocyclooctyne (DBCO) conjugated LD655 dye (Lumidyne Technologies) at 4 °C overnight. The reaction was quenched with 1 mM free AzF, followed by addition of 5 mM DTT to reverse the DNTB modification of cysteines. Finally, labeled base subcomplexes were purified using size-exclusion chromatography with a Superose 6 Increase 10/300 column (Cytiva) equilibrated in GF buffer (30 mM HEPES, pH 7.6, 50 mM NaCl, 50 mM KCl, 10 mM MgCl_2_, 5% glycerol, 1 mM ATP, 0.5 mM TCEP). Base concentrations were determined by Bradford assay, and labeling efficiencies were estimated by measuring the absorbance of the fluorophore.

Recombinant yeast lid was expressed from three plasmids, pAM80, pAM85, and pAM86 that code for Rpn3, Rpn5 – Rpn9, Rpn11, Rpn12, Sem1, and rare tRNAs, purified using a three-step procedure, and fluorescently labeled, as previously described ^4,53^. Briefly, fully assembled complexes containing His-Rpn12 and MBP-Rpn6 were purified using a HisTrap and amylose resin (NEB), and cleaved with HRV-protease. Subsequent fluorophore labeling and purification by size-exclusion chromatography proceeded in a similar fashion as for the base subcomplex.

The ubiquitin receptor subunit Rpn10 was expressed and purified separately as previously described ^4^. Pelleted cells were resuspended in lysis buffer supplemented with 20 mM imidazole, lysozyme, benzonase, and protease inhibitors, and lysed by sonication. After clarification, the lysate was batch bound to Ni-NTA affinity resin (ThermoFisher), the resin was washed with lysis buffer, and proteins were eluted with lysis buffer containing 250 mM imidazole. Rpn10 was further purified by size-exclusion chromatography using a Superdex 75 16/60 column (Cytiva). The ubiquitin receptor subunit Rpn13 was expressed and purified similar to Rpn10, with the added step of incubation with HRV-protease at 25 °C for 1 hour to remove a cleavable histidine tag before size exclusion chromatography.

### Cloning and purification of 20S core particle

A sequence encoding for an AviTag followed by a 3xFLAG tag was cloned into the N-terminus of the endogenous Pre1 gene of a W303 yeast strain. The purification of AviTag-20S core was performed similar to a procedure previously described ^53^. Briefly, the yeast strain was grown in YPD at 30 °C for 3 days. The cells were pelleted, resuspended in lysis buffer (60 mM HEPES, pH 7.6, 500 mM NaCl, 1 mM EDTA, 0.2% NP-40), popcorned into liquid nitrogen, and then lysed in a Cryomill 6875D (SPEXSamplePrep). The lysate was allowed to return to room temperature and clarified by centrifugation, and the 20S core particle was purified using anti-Flag M2 affinity resin (Sigma). The elution was concentrated using an Amicon Ultracel-100K, and the concentration was determined by absorbance at 280 nm. After biotinylation through incubation with 25 μM BirA and 100 μM biotin in the presence of 10 mM ATP and 10mM MgCl_2_ overnight at 4 °C, the core was further purified by size-exclusion chromatography with a Superose 6 Increase 10/300 column (Cytiva) equilibrated in GF buffer (30 mM HEPES, pH 7.6, 50 mM NaCl, 50 mM KCl, 10 mM MgCl_2_, 5% glycerol, 1 mM ATP, 0.5 mM TCEP). The extent of biotinylation was determined in a gel-shift assay by incubation with NeutrAvidin (ThermoFisher).

### Substrate purification and labeling

Titin I27 substrate expression was performed as described previously ^4^, using a plasmid that codes for the desired protein sequence fused to an N-terminal intein and chitin binding domain (CBD) as part of the IMPACT purification system (NEB). 2 liters of cells were grown to an OD_600_ of 0.6 and induced with 1 mM IPTG for 3 hours at 25 °C. Cells were pelleted, resuspended in chitin binding buffer (60 mM HEPES, pH 7.6, 150 mM NaCl, 1 mM EDTA, 5% glycerol) with benzonase, and protease inhibitors, and lysed. After lysate clarification by centrifugation, the protein was batch bound to 10 mL of chitin resin (NEB) for 1 hour at 4 °C. The resin was washed with 100 mL of chitin binding buffer, resuspended in cleavage buffer (60 mM HEPES, pH 8.5, 150 mM NaCl, 1 mM EDTA, 5% glycerol, and 50 mM DTT), and incubated overnight at 4 °C. The flowthrough containing the cleaved protein was collected, run over fresh chitin resin to remove any uncleaved protein, concentrated, and further purified using a Superdex 75 16/60 column (Cytiva) equilibrated with GF buffer. Fluorophores were covalently attached either to the substrates’ N-terminus for bulk degradation measurements or to single engineered cysteines for FRET-based single-molecule experiments. For N-terminal labeling with fluorescein, we used sortase A ^4,54^ and the peptide FAM-HHHHHHLPETGG (Genscript). For the labeling of engineered cysteines by maleimide chemistry, substrates were dialyzed into labeling buffer (30mM HEPES 7.2, 150 mM NaCl, 1 mM EDTA) for 3 hours at room temperature using Slide-a-Lyzer mini dialysis cups (ThermoFisher), diluted to 100 μM, and then incubated for 1 hour at room temperature with 200 μM fluorophore. Excess dye was quenched with 1 mM DTT and removed by size-exclusion chromatography on a Superdex 75 10/300 column equilibrated with GF buffer.

### Ubiquitin purification, substrate ubiquitination, and preparation of unanchored ubiquitin chains

*E. Coli* Bl21*(DE3) cells were transformed with a pET28a plasmid coding for yeast ubiquitin, grown in TB at 37 °C until OD_600_ = 0.6 - 0.9, induced with 1 mM IPTG overnight at 18 °C, pelleted, resuspended in lysis buffer with lysozyme, benzonase, and protease inhibitors, and lysed by sonication. The lysate was clarified by centrifugation and its pH adjusted to 4.5 using acetic acid. Protein precipitate was removed by centrifugation and the supernatant was dialyzed against 50 mM Na-acetate, pH 4.5 overnight at 4 °C. Ubiquitin was purified by cation-exchange chromatography with a 5 mL HiTrap SP FF column (Cytiva) and a gradient of 0 - 0.5 M NaCl in 50 mM Na-acetate, pH 4.5, followed by size-exclusion chromatography using a Superdex 75 16/60 column equilibrated with GF buffer. Enzymatic addition of ubiquitin chains to substrates containing PPPYX motifs and a single lysine residue was performed as previously described ^4^, incubating 10 μM substrate for 3 hours at 25 °C in GF buffer with 10 mM ATP, 400 μM ubiquitin, 2.5 μM mouse E1, 2.5 μM Ubc1, and 25 μM Rsp5. K48-linked tetra-ubiquitin chains were synthesized and purified as previously described ^55^: 1 mM ubiquitin was incubated with 1 μM mE1, 5 μM Cdc34, and 10 mM ATP in GF buffer overnight at 37 °C, and chains were separated using a Resource S cation-exchange column with an NaCl gradient. Tetra-ubiquitin chains were verified by gel electrophoresis.

### Microscope setup

Single-molecule data acquisition was performed at room temperature on a custom-built objective-type TIRF microscope, similar to a design described previously ^56^. The microscope is built on top of a Nikon Eclipse Ti turret that is equipped with an ultra-stable single-molecule stage (model KS-N) and a 60X Apo TIRF (1.49 N.A.) oil objective (Nikon), which we used with Cargille DF immersion oil. No active feedback stabilization was used, as the stage exhibited a drift of less than 0.5 pixels over 20 minutes (more than 10 times longer than a normal acquisition). Donor dyes (Cy3 and LD555) were imaged using a Coherent OBIS LS 100 mW 532 nm solid-state laser (operated at 2-10 mW power). Acceptor dyes (Cy5 and LD655) were imaged using a Uniphase 10 mW 633 nm helium-neon laser (operated at 10 mW and attenuated to ~5 mW using a N.D. filter). All lasers were steered through Galilean beam expanding optics to save space. The collimated beams were focused using an achromatic doublet (f = 100 mm) onto a conjugate back focal plane of the objective that was relayed outside of the microscope with a Ti-Lapp branch (Nikon). The focusing lens was coupled to a positioning stage and mirror assembly, such that the TIRF angle could be adjusted by moving the lens and mirror simultaneously. We used a filter cube from Chroma (TRF59907-EM). The emission from donor and acceptor dyes were separated using image splitting optics (W-View Gemini from Hamamatsu) and simultaneously imaged with the same camera (Andor iXon 512 × 512 EMCCD DU-897). For registration of the two emission channels, images were acquired with 100 nm TetraSpeck beads (ThermoFisher). For preparation of the TetraSpeck bead sample, poly-D-lysine (2 mg/ml) was flowed into a reaction chamber and incubated for 3 minutes, followed by a wash with GF buffer, before flowing in a 1:50 dilution of TetraSpeck beads and incubation for 5 min. Excess beads were washed out using GF buffer, and the sample was imaged using the 532 nm laser. Registration was then performed by overlaying cropped regions from the donor and acceptor emission channels such that the bead positions in each channel agreed. All acquisitions were performed with the following camera settings: EM Gain at 60, readout rate at 17 MHz, preamp set to 3, with activated overlap mode and camera internally triggered.

### Single-molecule fluorescence microscopy data acquisition

Reaction chambers were assembled on microscope slides using double sided Scotch tape and PEG coated coverslips with low density PEG-biotin (Microsurfaces Inc.). The reaction chambers were incubated with 0.05 mg/ml NeutrAvidin (ThermoFisher), and excess NeutrAvidin was removed by washing with 50 μL assay buffer (GF buffer supplemented with 0.3 μM Rpn10, 1.2 mM Trolox, 0.2 mM beta-mercaptoethanol, 2 mM ATP, and an ATP-regeneration system consisting of 0.03 mg/mL creatine kinase and 16 mM creatine phosphate). The assay buffer for the conformational change assay during substrate processing was further supplemented with 0.3 μM Rpn13 to assure complete proteasome assemblies and because this subunit was previously reported to be required for unfoldase activation ^57^. A dilution of dye-labeled proteasomes (around ~100 pM) was flowed into the reaction chamber and allowed to incubate for 5 minutes. The extent of surface deposition was controlled by adjusting the dilution of incubated proteasomes, such that single fluorescent molecules could clearly be distinguished. Excess proteasomes in solution were washed out with 50 μL assay buffer followed by 50 μL Imaging Buffer (assay buffer supplemented with the PCA/PCD oxygen scavenging system described previously ^58^). Substrate was added to the Imaging Buffer for assays that required it. For conformational change assay, saturating amount of substrate was added at a concentration of either 400 nM for ubiquitinated substrate and 5 μM for SspB delivered substrate. For the substrate-processing assay, donor labeled substrate was added at a concentration of 25 nM.

### Data analysis

For the conformational dynamics measurements, raw images were cropped and spatially aligned using registration coordinates derived from analyzing TetraSpeck images. These movies were then parsed by the Spartan software package ^59^. Once parsed, the traces were sorted, analyzed, and plotted using Spartan’s built-in functions. Raw images from substrate processing acquisitions were similarly registered and then parsed using the ImageJ plugin spotIntensityInAllChannels ^60^. The parsed data were sorted and analyzed using a custom-built Matlab app. To calculate FRET efficiency, the raw donor and acceptor fluorescence traces were smoothed using a moving average filter – Matlab’s smooth function with default settings. Acceptor fluorescence was then corrected for donor bleedthrough used to calculate “filtered FRET”. Individual traces were then inspected manually and scored for various features. The start and end of each reaction was determined from the total fluorescence, which is determined by the total residency time of a donor labeled substrate. The tail-insertion phase was measured from the appearance of a donor signal and intermediate FRET to the point of the first high FRET peak. The FRET decay was scored similarly but in reverse, i.e. by measuring the time from a high FRET peak to the point where the signal decayed to below a threshold of 0.2. The length of the high-FRET phase was determined as the time between the tail insertion (FRET-increase) phase and the FRET-decay phase. Integration time for these measurements was 100 ms, which also sets a limit for the processes we can distinguish.

Substrate-processing dwells in the conformational dynamics assay were scored manually from the onset of high FRET to the stable return to low FRET, including brief excursions to low FRET during the processing of the I27^V15P^ and I27^WT^ substrates. The distributions of each measurement (tail insertion, processing dwells, etc.) were fit to a gamma distribution using Matlab’s fitdist function. The parameters of the fit were then used to generate fit lines for 1-cumulative distribution functions (1-CDF) using Matlab’s gamcdf function. Empirically derived 1-CDF plots were generated from the measurements using Matlab’s ecdf function. The conformational change assays were performed at a sampling rate of 20 Hz and analyzed using Hidden Markov modeling with ebFRET ^61^. The dwell time distributions for each FRET state were extracted and fit to a single exponential, with errors representing 95% confidence interval of the fit.

To determine the substrate-capture success of the proteasome, we calculated the fractional number of productive binding events relative to the total number of substrate-binding events (both productive and non-productive). We scored as binding events all events that exhibited an increase in donor fluorescence colocalized with acceptor fluorescence. Productive binding were considered those events that exhibited a donor-fluorescence recovery after a high-FRET phase as a measure of successful threading. Non-productive binding often appeared as intermediate FRET without an increase to high FRET or recovery of the donor fluorescence. Five movies, containing at least 100 binding events per movie, were analyzed for each condition, and the error is the standard deviation for the five measurements.

### Fluorescence polarization-based multiple-turnover degradation assay

Proteasomes were reconstituted at 2x concentration by mixing and incubating 100 nM core particle, 800 nM SspB-fused base, 1.2 μM lid and 2 μM Rpn10 at room temperature for 10 min in the presence of an ATP regeneration system (0.03 mg/mL creatine kinase and 16 mM creatine phosphate). Substrate mix was prepared at 2x concentration (20 μM) in GF buffer supplemented with 1 mg/mL BSA and 10 mM ATP. For the experiments with K48-linked tetra-ubiquitin chains, 20 μM chains were added in the substrate mix. The multiple-turnover degradation reaction was initiated by mixing 7 μL of 2x reconstituted SspB-fused proteasome and 7 μL of 2x substrate mix. 10 μL of the reaction was immediately transferred to a 384-well low-volume black flat bottom plate (Corning #3820) pre-warmed to 30 °C, and degradation was monitored by the loss of fluorescence-polarization signal from 5-carboxyfluorescein-labeled substrates in a CLARIOstarPlus plate reader (BMG Labtech) at 30 °C. Rates for multiple-turnover degradation were determined by linear regression of the initial change in polarization and normalizing the fit with the measured differences in fluorescence-polarization signals of undegraded substrates and substrates fully degraded by 0.1 μg/μL chymotrypsin.

## Data Availability

All data generated or analyzed during this study are included in this manuscript and the supplementary information.

## Code Availability

Custom code for single-molecule data analysis will be available upon request.

## REFERENCES

1 Chen, B., Retzlaff, M., Roos, T. & Frydman, J. Cellular strategies of protein quality control. Cold Spring Harb Perspect Biol 3, a004374, doi:10.1101/cshperspect.a004374 (2011).

2 Hershko, A. & Ciechanover, A. The ubiquitin system. Annu Rev Biochem 67, 425–479, doi:10.1146/annurev.biochem.67.1.425 (1998).

3 Bard, J. A. M. et al. Structure and Function of the 26S Proteasome. Annu Rev Biochem 87, 697–724, doi:10.1146/annurev-biochem-062917-011931 (2018).

4 Bard, J. A. M., Bashore, C., Dong, K. C. & Martin, A. The 26S Proteasome Utilizes a Kinetic Gateway to Prioritize Substrate Degradation. Cell 177, 286–298 e215, doi:10.1016/j.cell.2019.02.031 (2019).

5 Prakash, S., Tian, L., Ratliff, K. S., Lehotzky, R. E. & Matouschek, A. An unstructured initiation site is required for efficient proteasome-mediated degradation. Nat Struct Mol Biol 11, 830–837, doi:10.1038/nsmb814 (2004).

6 Fishbain, S., Prakash, S., Herrig, A., Elsasser, S. & Matouschek, A. Rad23 escapes degradation because it lacks a proteasome initiation region. Nature communications 2, 192, doi:10.1038/ncomms1194 (2011).

7 Inobe, T., Fishbain, S., Prakash, S. & Matouschek, A. Defining the geometry of the two-component proteasome degron. Nat. Chem. Biol. 7, 161–167, doi:10.1038/nchembio.521 (2011).

8 Fishbain, S. et al. Sequence composition of disordered regions fine-tunes protein half-life. Nature Structural & Molecular Biology 22, 214–221, doi:10.1038/nsmb.2958 (2015).

9 Takeuchi, J., Chen, H. & Coffino, P. Proteasome substrate degradation requires association plus extended peptide. The EMBO Journal 26, 123–131, doi:10.1038/sj.emboj.7601476 (2007).

10 Peth, A., Uchiki, T. & Goldberg, A. L. ATP-dependent steps in the binding of ubiquitin conjugates to the 26S proteasome that commit to degradation. Mol. Cell 40, 671–681, doi:10.1016/j.molcel.2010.11.002 10.1016/j.molcel.2010.11.002. (2010).

11 Groll, M. et al. A gated channel into the proteasome core particle. Nat Struct Biol 7, 1062–1067 (2000).

12 Matyskiela, M. E. & Martin, A. Design principles of a universal protein degradation machine. J Mol Biol 425, 199–213, doi:10.1016/j.jmb.2012.11.001 10.1016/j.jmb.2012.11.001. Epub 2012 Nov 9. (2013).

13 Saeki, Y. & Tanaka, K. Assembly and function of the proteasome. Methods Mol Biol 832, 315–337, doi:10.1007/978-1-61779-474-2_22 10.1007/978-1-61779-474-2_22. (2012).

14 Shi, Y. et al. Rpn1 provides adjacent receptor sites for substrate binding and deubiquitination by the proteasome. Science 351, aad9421, doi:10.1126/science.aad9421 (2016).

15 Deveraux, Q., Ustrell, V., Pickart, C. & Rechsteiner, M. A 26 S protease subunit that binds ubiquitin conjugates. J. Biol. Chem. 269, 7059–7061 (1994).

16 Husnjak, K. et al. Proteasome subunit Rpn13 is a novel ubiquitin receptor. Nature 453, 481–488, doi:10.1038/nature06926 (2008).

17 Glickman, M. H., Rubin, D. M., Fried, V. A. & Finley, D. The regulatory particle of the Saccharomyces cerevisiae proteasome. Molecular and cellular biology 18, 3149–3162 (1998).

18 Tomko, R. J. Jr & Hochstrasser, M. Molecular Architecture and Assembly of the Eukaryotic Proteasome. Annu Rev Biochem, doi:10.1146/annurev-biochem-060410-150257 (2013).

19 Tomko, R. J., Jr., Funakoshi, M., Schneider, K., Wang, J. & Hochstrasser, M. Heterohexameric ring arrangement of the eukaryotic proteasomal ATPases: implications for proteasome structure and assembly. Mol Cell 38, 393–403, doi:10.1016/j.molcel.2010.02.035 (2010).

20 Erales, J., Hoyt, M. A., Troll, F. & Coffino, P. Functional asymmetries of proteasome translocase pore. J Biol Chem 287, 18535–18543, doi:10.1074/jbc.M112.357327 (2012).

21 Beckwith, R., Estrin, E., Worden, E. J. & Martin, A. Reconstitution of the 26S proteasome reveals functional asymmetries in its AAA+ unfoldase. Nat Struct Mol Biol 20, 1164–1172, doi:10.1038/nsmb.2659 10.1038/nsmb.2659. Epub 2013 Sep 8. (2013).

22 Verma, R. et al. Role of Rpn11 metalloprotease in deubiquitination and degradation by the 26S proteasome. Science 298, 611–615 (2002).

23 Yao, T. & Cohen, R. E. A cryptic protease couples deubiquitination and degradation by the proteasome. Nature 419, 403–407, doi:10.1038/nature01071 (2002).

24 Worden, E. J., Padovani, C. & Martin, A. Structure of the Rpn11-Rpn8 dimer reveals mechanisms of substrate deubiquitination during proteasomal degradation. Nature Structural & Molecular Biology 21, 220–227, doi:10.1038/nsmb.2771 10.1038/nsmb.2771. Epub 2014 Jan 23. (2014).

25 Pathare, G. R. et al. Crystal structure of the proteasomal deubiquitylation module Rpn8-Rpn11. Proc Natl Acad Sci USA 111, 2984–2989, doi:10.1073/pnas.1400546111 10.1073/pnas.1400546111. Epub 2014 Feb 10. (2014).

26 Lander, G. C. et al. Complete subunit architecture of the proteasome regulatory particle. Nature 482, 186–191, doi:10.1038/nature10774 10.1038/nature10774. (2012).

27 Worden, E. J., Dong, K. C. & Martin, A. An AAA Motor-Driven Mechanical Switch in Rpn11 Controls Deubiquitination at the 26S Proteasome. Mol. Cell 67, 799–811.e798, doi:10.1016/j.molcel.2017.07.023 10.1016/j.molcel.2017.07.023. (2017).

28 Matyskiela, M. E., Lander, G. C. & Martin, A. Conformational switching of the 26S proteasome enables substrate degradation. Nat Struct Mol Biol 20, 781–788, doi:10.1038/nsmb.2616 10.1038/nsmb.2616. Epub 2013 Jun 16. (2013).

29 Sledz, P. et al. Structure of the 26S proteasome with ATP-gammaS bound provides insights into the mechanism of nucleotide-dependent substrate translocation. Proc Natl Acad Sci USA 110, 7264–7269, doi:10.1073/pnas.1305782110 10.1073/pnas.1305782110. Epub 2013 Apr 15. (2013).

30 Unverdorben, P. et al. Deep classification of a large cryo-EM dataset defines the conformational landscape of the 26S proteasome. Proc Natl Acad Sci USA 111, 5544–5549, doi:10.1073/pnas.1403409111 10.1073/pnas.1403409111. Epub 2014 Mar 24. (2014).

31 Asano, S. et al. A molecular census of 26S proteasomes in intact neurons. Science 347, 439–442 (2015).

32 Wehmer, M. et al. Structural insights into the functional cycle of the ATPase module of the 26S proteasome. Proc Natl Acad Sci USA 114, 1305–1310, doi:10.1073/pnas.1621129114 10.1073/pnas.1621129114. Epub 2017 Jan 23. (2017).

33 Eisele, M. R. et al. Expanded Coverage of the 26S Proteasome Conformational Landscape Reveals Mechanisms of Peptidase Gating. Cell Rep 24, 1301–1315 e1305, doi:10.1016/j.celrep.2018.07.004 (2018).

34 Ding, Z. et al. High-resolution cryo-EM structure of the proteasome in complex with ADP-AlFx. Cell Res 27, 373–385, doi:10.1038/cr.2017.12 10.1038/cr.2017.12. Epub 2017 Jan 20. (2017).

35 de la Pena, A. H., Goodall, E. A., Gates, S. N., Lander, G. C. & Martin, A. Substrate-engaged 26S proteasome structures reveal mechanisms for ATP-hydrolysis-driven translocation. Science 362, doi:10.1126/science.aav0725 (2018).

36 Dong, Y. et al. Cryo-EM structures and dynamics of substrate-engaged human 26S proteasome. Nature, doi:10.1038/s41586-018-0736-4 (2018).

37 Puchades, C. et al. Structure of the mitochondrial inner membrane AAA+ protease YME1 gives insight into substrate processing. Science 358, doi:10.1126/science.aao0464 (2017).

38 Han, H., Monroe, N., Sundquist, W. I., Shen, P. S. & Hill, C. P. The AAA ATPase Vps4 binds ESCRT-III substrates through a repeating array of dipeptide-binding pockets. eLife 6, doi:10.7554/eLife.31324 (2017).

39 Monroe, N., Han, H., Shen, P. S., Sundquist, W. I. & Hill, C. P. Structural basis of protein translocation by the Vps4-Vta1 AAA ATPase. eLife 6, doi:10.7554/eLife.24487 (2017).

40 Greene, E. R. et al. Specific lid-base contacts in the 26s proteasome control the conformational switching required for substrate degradation. eLife 8, doi:10.7554/eLife.49806 (2019).

41 Nemec, A. A., Peterson, A. K., Warnock, J. L., Reed, R. G. & Tomko, R. J., Jr. An Allosteric Interaction Network Promotes Conformation State-Dependent Eviction of the Nas6 Assembly Chaperone from Nascent 26S Proteasomes. Cell Rep 26, 483–495 e485, doi:10.1016/j.celrep.2018.12.042 (2019).

42 Reichard, E. L. et al. Substrate Ubiquitination Controls the Unfolding Ability of the Proteasome. J Biol Chem 291, 18547–18561, doi:10.1074/jbc.M116.720151 (2016).

43 Braganca, C. E. & Kraut, D. A. Mode of targeting to the proteasome determines GFP fate. J Biol Chem 295, 15892–15901, doi:10.1074/jbc.RA120.015235 (2020).

44 Kenniston, J. A., Baker, T. A., Fernandez, J. M. & Sauer, R. T. Linkage between ATP consumption and mechanical unfolding during the protein processing reactions of an AAA+ degradation machine. Cell 114, 511–520 (2003).

45 Maillard, R. A. et al. ClpX(P) generates mechanical force to unfold and translocate its protein substrates. Cell 145, 459–469, doi:10.1016/j.cell.2011.04.010 (2011).

46 Sen, M. et al. The ClpXP protease unfolds substrates using a constant rate of pulling but different gears. Cell 155, 636–646, doi:10.1016/j.cell.2013.09.022 10.1016/j.cell.2013.09.022. Epub 2013 Oct 24. (2013).

47 Aubin-Tam, M. E., Olivares, A. O., Sauer, R. T., Baker, T. A. & Lang, M. J. Single-molecule protein unfolding and translocation by an ATP-fueled proteolytic machine. Cell 145, 257–267, doi:10.1016/j.cell.2011.03.036 (2011).

48 Olivares, A. O., Kotamarthi, H. C., Stein, B. J., Sauer, R. T. & Baker, T. A. Effect of directional pulling on mechanical protein degradation by ATP-dependent proteolytic machines. Proc Natl Acad Sci USA 114, E6306–E6313, doi:10.1073/pnas.1707794114 (2017).

49 Sakata, E. et al. Localization of the proteasomal ubiquitin receptors Rpn10 and Rpn13 by electron cryomicroscopy. Proc. Natl. Acad. Sci. U.S.A. 109, 1479–1484, doi:10.1073/pnas.1119394109 (2012).

50 Bashore, C. et al. Ubp6 deubiquitinase controls conformational dynamics and substrate degradation of the 26S proteasome. Nature structural & molecular biology 22, 712–719, doi:10.1038/nsmb.3075 10.1038/nsmb.3075. Epub 2015 Aug 24. (2015).

51 Hersch, G. L., Baker, T. A. & Sauer, R. T. SspB delivery of substrates for ClpXP proteolysis probed by the design of improved degradation tags. Proc Natl Acad Sci USA 101, 12136–12141, doi:10.1073/pnas.0404733101 (2004).

52 Cresti, J. R. et al. Proteasomal conformation controls unfolding ability. Proc Natl Acad Sci USA 118, doi:10.1073/pnas.2101004118 (2021).

53 Bard, J. A. M. & Martin, A. Recombinant Expression, Unnatural Amino Acid Incorporation, and Site-Specific Labeling of 26S Proteasomal Subcomplexes. Methods Mol Biol 1844, 219–236, doi:10.1007/978-1-4939-8706-1_15 (2018).

54 Theile, C. S. et al. Site-specific N-terminal labeling of proteins using sortase-mediated reactions. Nat Protoc 8, 1800–1807, doi:10.1038/nprot.2013.102 (2013).

55 Dong, K. C. et al. Preparation of distinct ubiquitin chain reagents of high purity and yield. Structure 19, 1053–1063, doi:10.1016/j.str.2011.06.010 10.1016/j.str.2011.06.010. (2011).

56 Niekamp, S. et al. Nanometer-accuracy distance measurements between fluorophores at the single-molecule level. Proc Natl Acad Sci USA 116, 4275–4284, doi:10.1073/pnas.1815826116 (2019).

57 Cundiff, M. D. et al. Ubiquitin receptors are required for substrate-mediated activation of the proteasome’s unfolding ability. Sci Rep 9, 14506, doi:10.1038/s41598-019-50857-y (2019).

58 Aitken, C. E., Marshall, R. A. & Puglisi, J. D. An oxygen scavenging system for improvement of dye stability in single-molecule fluorescence experiments. Biophys J 94, 1826–1835, doi:10.1529/biophysj.107.117689 (2008).

59 Juette, M. F. et al. Single-molecule imaging of non-equilibrium molecular ensembles on the millisecond timescale. Nat Methods 13, 341–344, doi:10.1038/nmeth.3769 (2016).

60 Schneider, C. A., Rasband, W. S. & Eliceiri, K. W. NIH Image to ImageJ: 25 years of image analysis. Nat Methods 9, 671–675, doi:10.1038/nmeth.2089 (2012).

61 van de Meent, J. W., Bronson, J. E., Wiggins, C. H. & Gonzalez, R. L., Jr. Empirical Bayes methods enable advanced population-level analyses of single-molecule FRET experiments. Biophys J 106, 1327–1337, doi:10.1016/j.bpj.2013.12.055 (2014).

