## Supplementary Figures and Tables for "Ubiquitin modulates 26S proteasome conformational dynamics and promotes substrate degradation"

**a** Heterologous expression and reconstitution system

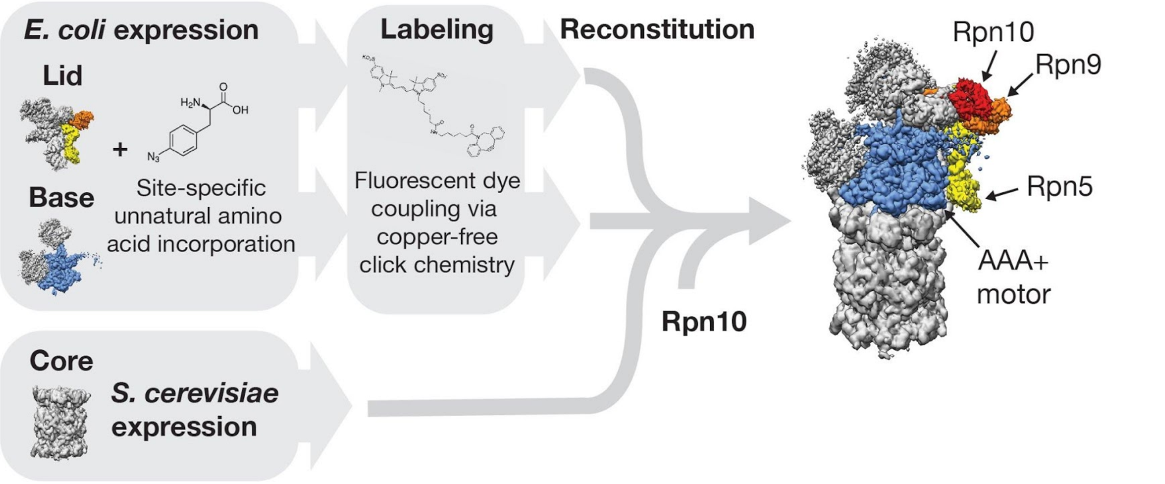


| Donor position | Acceptor position | Protasome conformation | PDB ID | Donor-acceptor distance (Å) |
|---|---|---|---|---|
| Rpn9 F2AzF | Rpt5 Q49AzF | s1 | 5MP9 | 77 |
| Rpn9 F2AzF | Rpt5 Q49AzF | non-s1 (s3) | 5MPB | 41 |
| I27-tail C127 | Rpt1 I191AzF | non-s1/substrate engaged | 6EF3 | 23 |

**b** Substrate-protein design

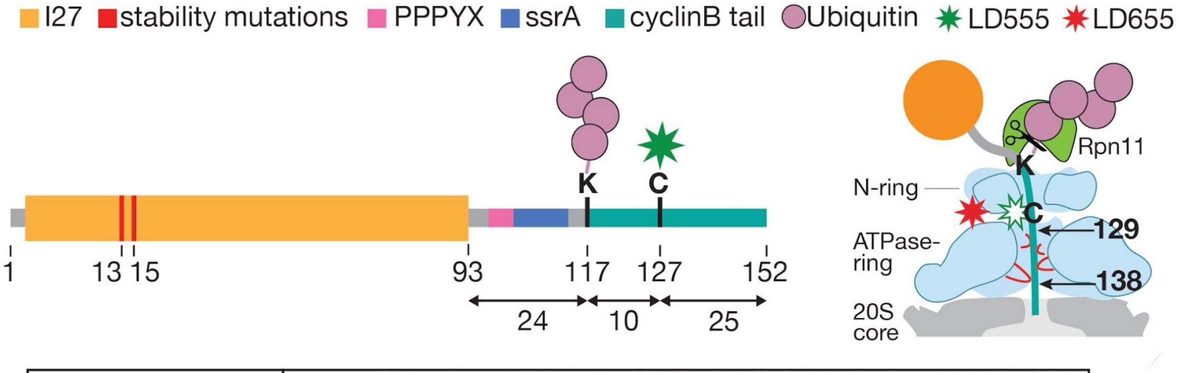


| | |
|---|---|
| I27 substrate sequence | MGGGLIEVERPLYGVEVFVGETAHFEIELSEPDVHGQWRLRGQPLAASPDCEIIEDGRRHILILHNCQLGMTGEVSFQAANTRSAANLRVRELGGAGPPPYSAANDENYALAAHGGKHTFNNENVSCRLGGAASIAVQAPAQHTFNNENVSY |

**Supplementary Figure 1: Heterologous proteasome expression and substrate-protein design. a**) Proteasome lid and base subcomplexes from *Saccharomyces cerevisiae* are heterologously expressed in *E. coli*, site-specifically labeled through incorporation of the unnatural amino acid azido-phenylalanine (AzF) and copper-free click addition of fluorescent dyes, and reconstituted with 20S core particle purified from *S. cerevisiae*. The table lists the labeling positions, the considered proteasome structures and conformational states, and the Cα distances between labeled residues for the FRET-based substrate-processing and conformational-change assays. **b**) Substrate proteins include the titin I27 domain from which all lysines have been removed and a C-terminally fused unstructured initiation region or "tail" derived from a Cyclin-B sequence. The tail contains a single lysine and PPPY motif for Rsp5-catalyzed polyubiquitination, a single cysteine for maleimide labeling with a donor fluorophore (LD555 or Cy3), and an ssrA sequence for binding to proteasome-fused SspB in the ubiquitin-independent delivery system. A N-terminal GGG motif was used to attach of fluorescein (FAM) in a sortase-catalyzed reaction for substrate-degradation measurements in bulk. The thermodynamic stability of the I27 domain can be modulated by incorporation of the V13P and V15P point mutations. The schematic on the right is based on our previous cryo-EM structure [35] and shows the substrate-engaged state of the proteasome prior to deubiquitination (PDB ID: 6EF3). For the substrate-processing assay, dye positions on the substrate and the proteasome were chosen to give a high-FRET signal in this engaged state after complete tail insertion, in which ~ 138 residues of the I27 model substrate reside above or inside the central channel of the motor.

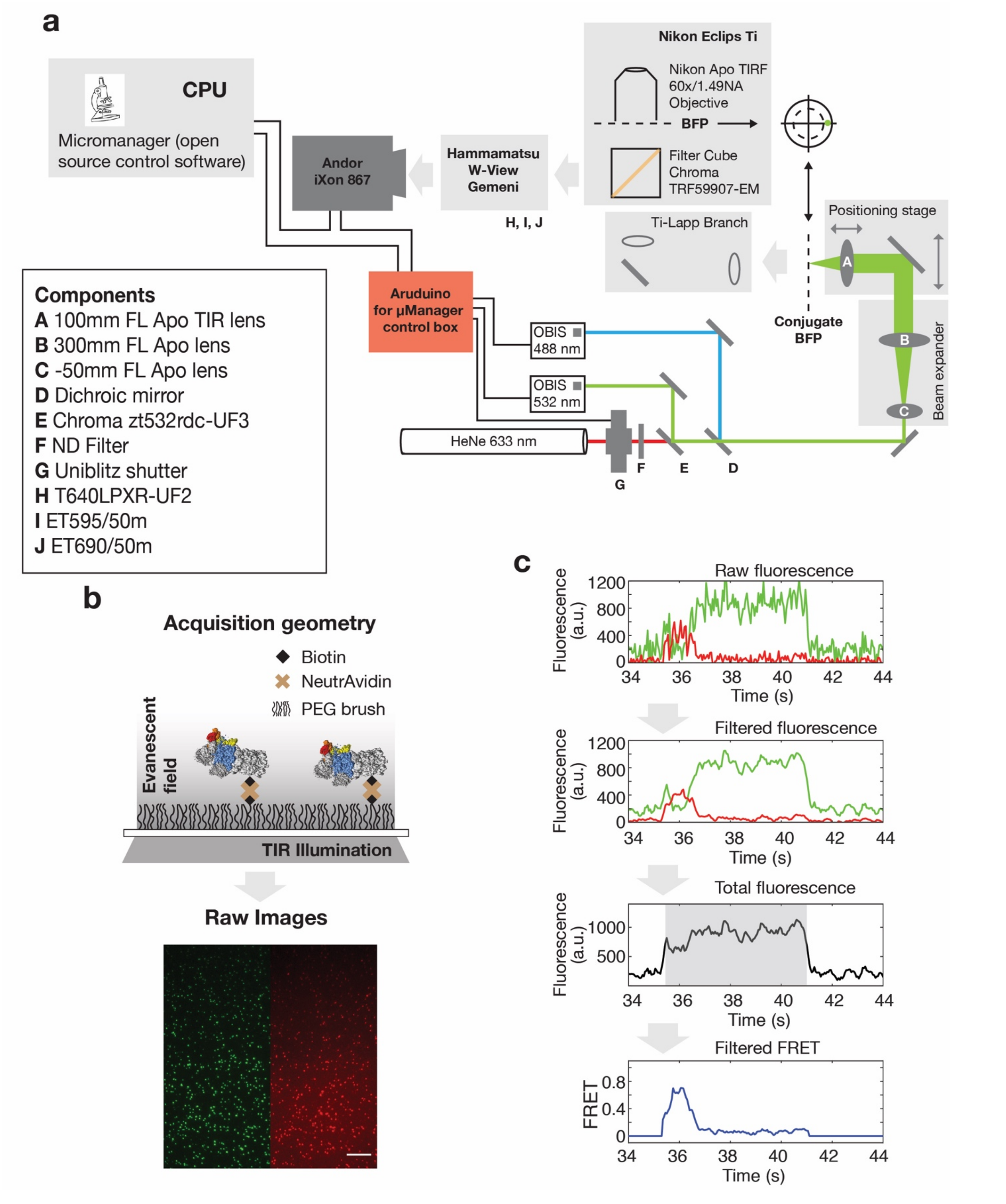


**Supplementary Figure 2: Design of the objective-type total internal reflection fluorescence (TIRF) microscope. a)** Diagram of the custom-built TIRF microscope used in this study. **b)** Cartoon representation (top) depicting the assay geometry. Singly labeled proteasomes for the substrate-processing assay or doubly labeled proteasome for the conformational-change assay were immobilized to the surface of a cover glass through biotin-

NeutrAvidin interaction. Raw data (bottom) for the FRET-based conformational change assay with LD555 donor-labeled lid (green) and LD655 acceptor-labeled base (red). Scale bar represents 10 μm. **c)** For the substrate processing assay, raw fluorescence is extracted from the microscope images and processed for subsequent analyses. Total fluorescence is used to determine the beginning and end of a substrate degradation event, as depicted by the grey shading. Filtered FRET is used to analyze the kinetics of individual processing steps.

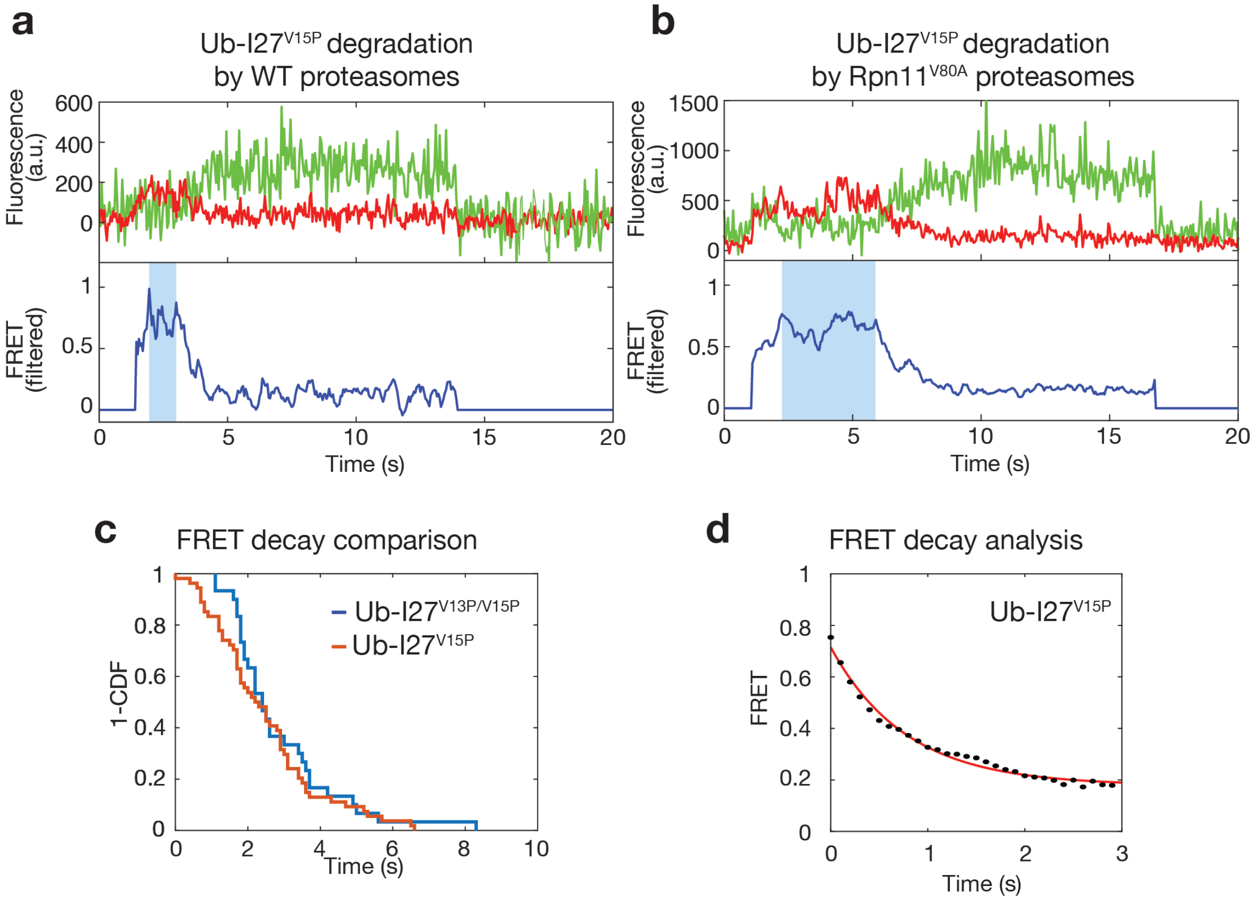


**Supplementary Figure 3: Representative traces for the substrate-processing assay. a)** Example trace for the degradation of ubiquitinated I27$^{V15P}$ substrate by wild-type proteasome, with the blue overlay indicating the high FRET phase. **b)** Example trace for the degradation of ubiquitinated I27$^{V15P}$ substrate by Rpn11$^{V80A}$-mutant proteasome, showing an extended high-FRET dwell (blue shading). **c)** Survival plots of the FRET-decay times for the degradation of ubiquitinated I27$^{V13P/V15P}$ and I27$^{V15P}$ substrates (N = 30 and 53, respectively). **d)** FRET-decay traces (N = 53) for the degradation of ubiquitinated I27$^{V15P}$ were averaged and fit to a single exponential (red line). The time constant derived from this fit is 0.78 ± 0.10 seconds (error is 95% confidence interval of the fit).

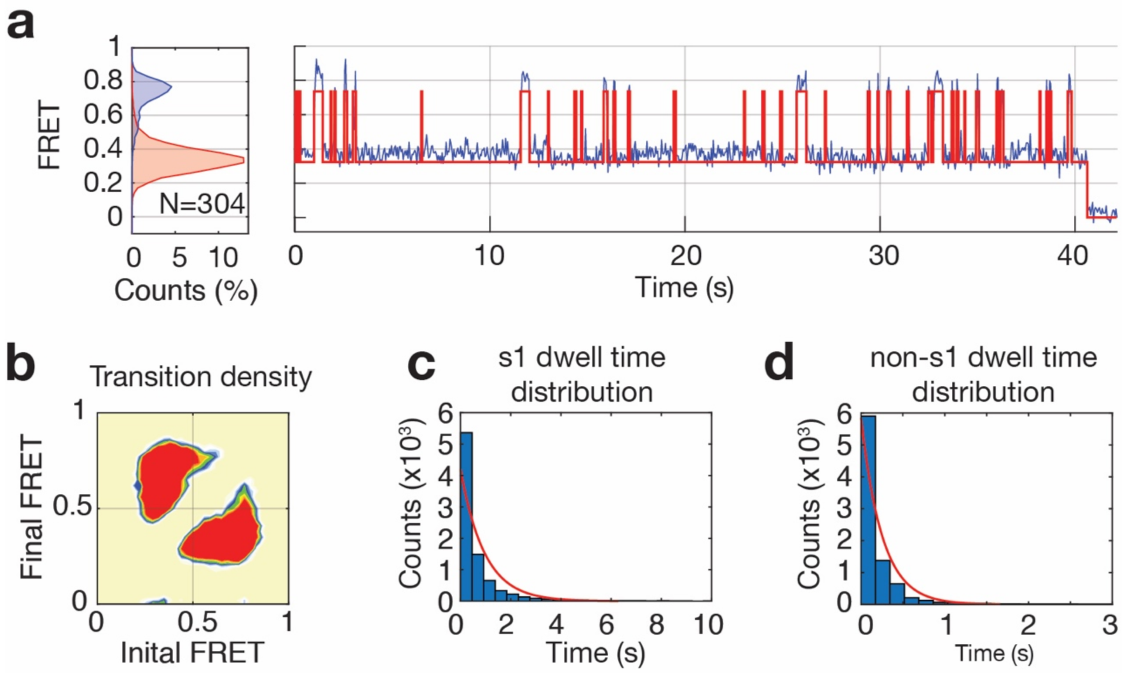


**Supplementary Figure 4: Hidden Markov modeling of conformational switching. a)** Representative trace of the conformational-change assay for wild-type proteasome in ATP, with dynamic switching fit according to a two-state system (red trace). The histogram on the left for multiple traces shows the relative occupancy of each state. **b)** Transition density plot depicting FRET values before and after each transition. **c)** Dwell-time distribution for the low-FRET s1 state is fit to a single exponential (red line) from which transition rates can be derived. **d**) Dwell-time distribution for the high-FRET non-s1 states is fit to a single exponential (red line).

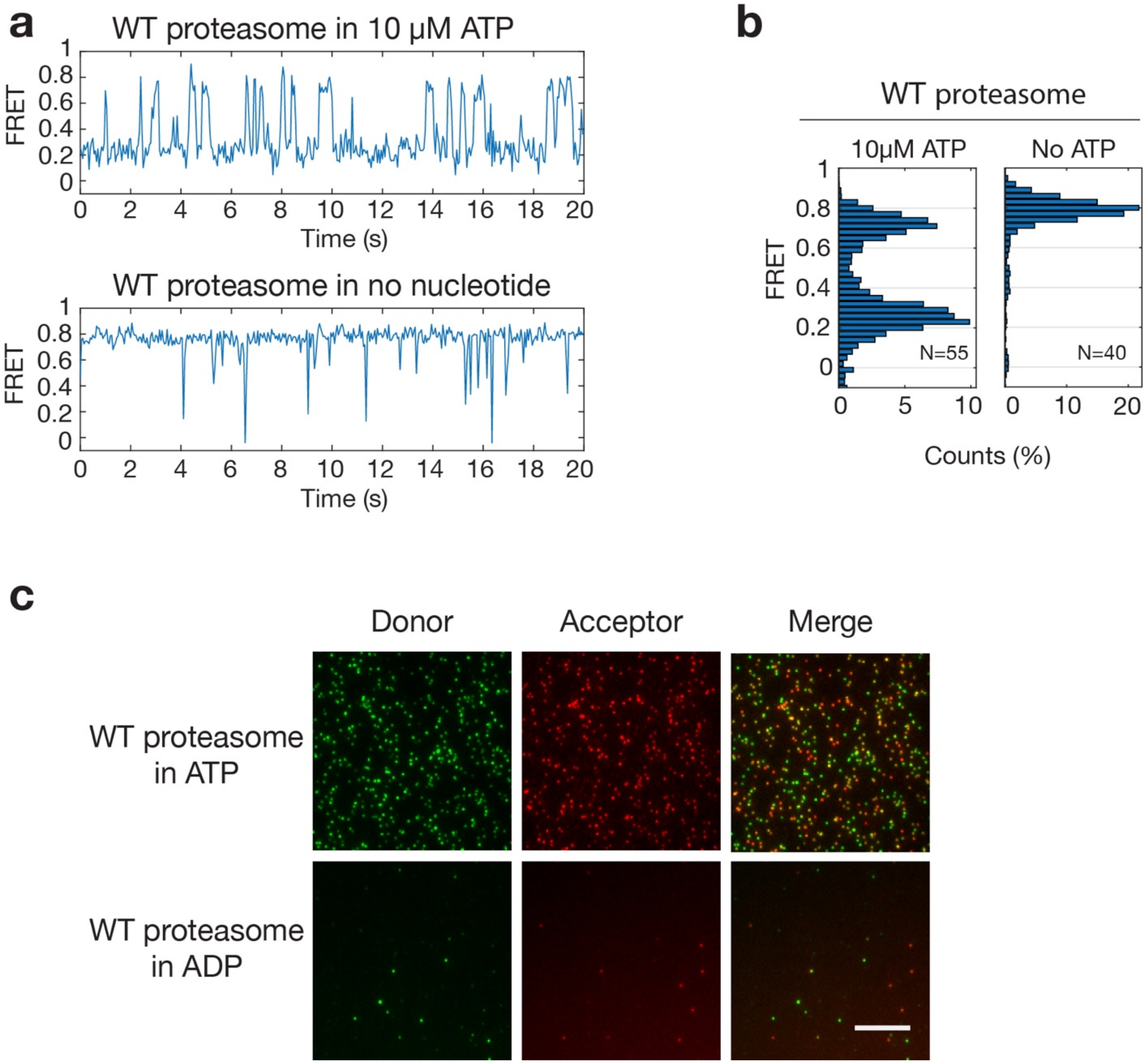


**Supplementary Figure 5: Nucleotide dependence of proteasome conformational dynamics. a)** Representative trace of wild-type proteasome in 10 μM ATP (top) exhibits more frequent and longer dwells for non-s1 states than at saturating ATP concentrations (c.f. Fig. 2b). Representative trace of wild-type proteasome in the absence of ATP (bottom) shows dominant non-s1 states with rare, brief transitions to the s1 state. **b)** Histograms for wild-type proteasomes at low ATP concentrations and without ATP show a bias towards the high-FRET non-s1 states. **c)** ADP promotes disassembly of the proteasome holoenzyme. Raw TIRF fluorescence images (excitation at 532 nm) of reconstituted proteasomes with donor-labeled lid (green) and acceptor-labeled base (red) show stable association at saturating ATP concentrations (top), but dissociation after buffer exchange with ADP (bottom). Scale bar represents 10 μm.

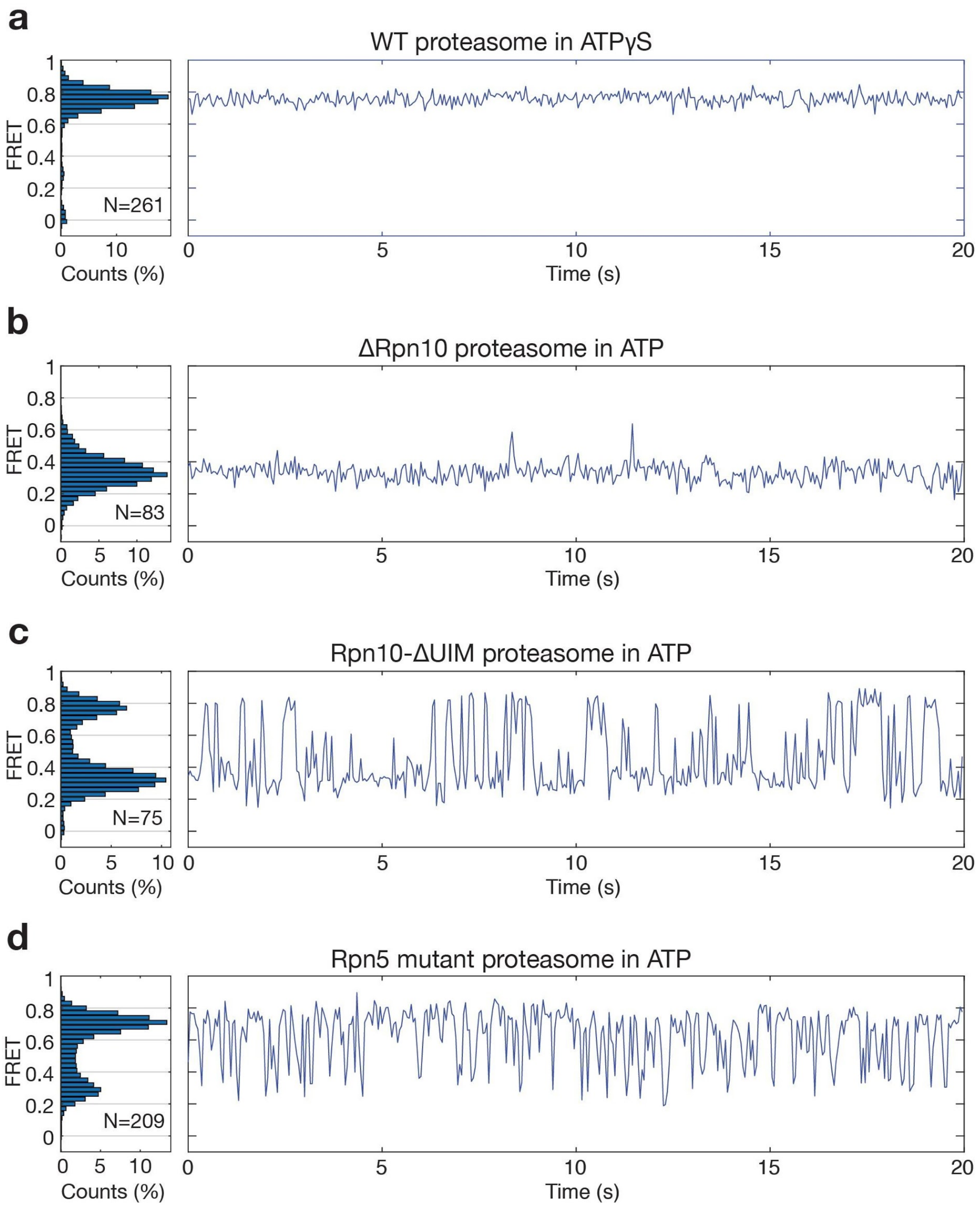


**Supplementary Figure 6: Representative conformational-change traces for proteasome under various conditions. a)** Example trace for the conformational changes of wild-type proteasome in ATPγS exhibits a stable high-FRET, non-s1 state. The histogram on the left indicates that the entire population is similarly biased towards non-s1 states. **b)** Example trace for the ΔRpn10 proteasome shows almost complete conformational bias towards the low FRET s1 state. A histogram for a population of ΔRpn10 proteasomes is depicted on the left. **c)** Representative trace for Rpn10-ΔUIM proteasomes exhibits longer dwells in high-FRET non-s1 states, consistent with the UIM stabilizing the s1 state (c.f. Supp. Table 2). The population histogram (left) for this mutant proteasome exhibits a more pronounced high-FRET compared to the wild-type proteasome (c.f. Figure 2b). **d)** Representative trace and histogram for Rpn5-mutant holoenzymes exhibits frequent switching and bias towards the high-FRET non-s1 states.

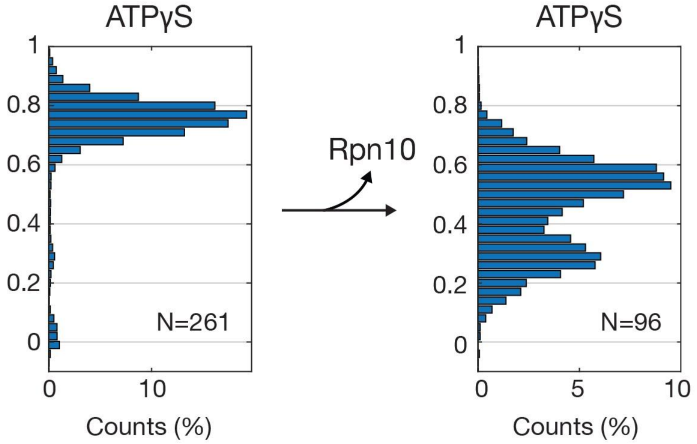


**Supplementary Figure 7: Rpn10 stabilizes the non-s1 state.** Histograms for the conformational equilibria of wild-type (left) and ΔRpn10 proteasomes (right) in ATPγS show a shift from a dominant non-s1 population to a bimodal distribution upon Rpn10 deletion. Non-s1 states of the ΔRpn10 proteasome show slightly lower FRET values (~ 0.6 versus ~ 0.8 for wild type), likely due to a reorientation of the Rpt5-attached acceptor dye.

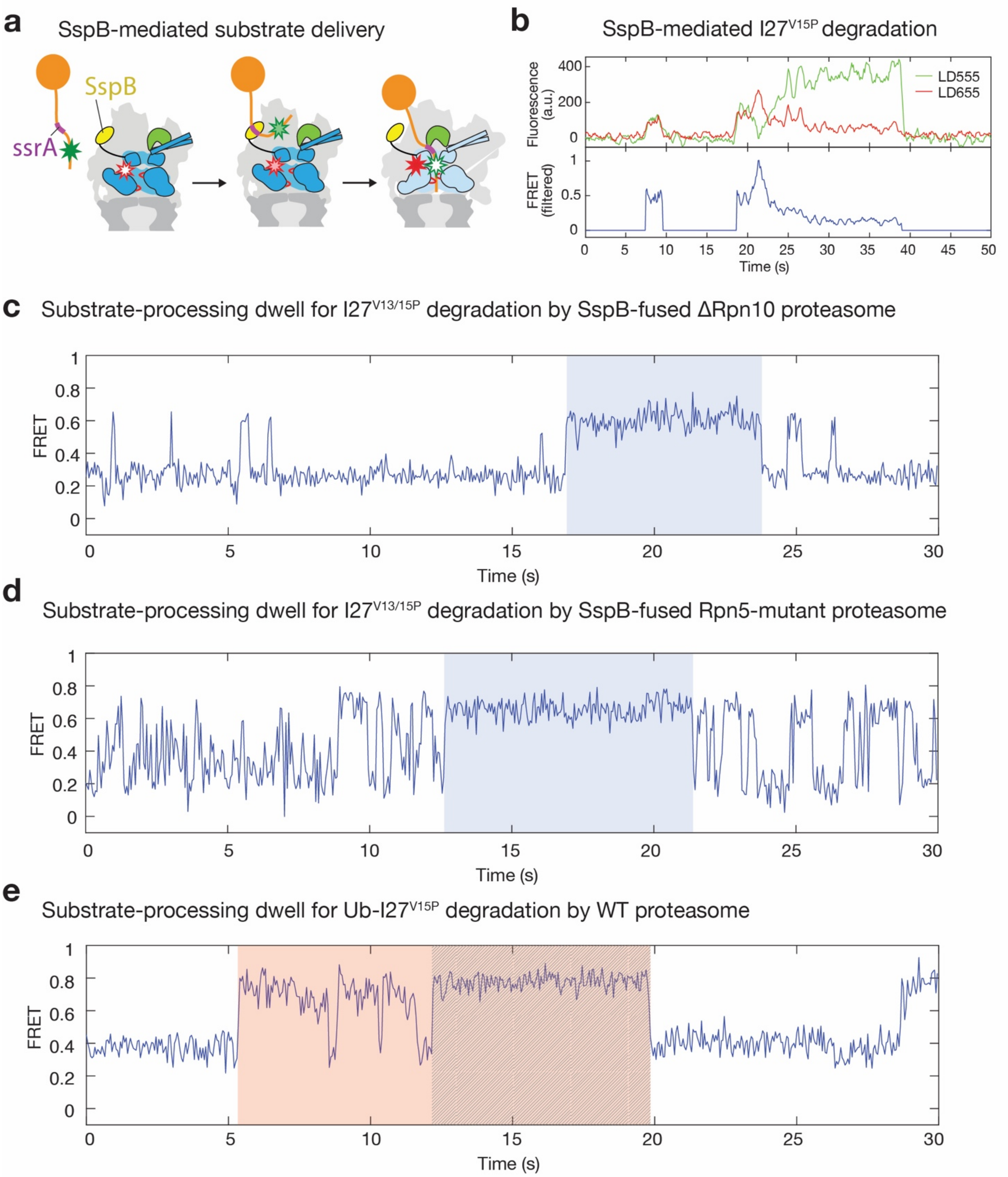


**Supplementary Figure 8: SspB-mediated substrate delivery and substrate-induced transitions to non-s1 states. a)** Schematic for ubiquitin-independent substrate delivery through the interaction between the proteasome-fused SspB (yellow) and the ssrA sequence (magenta) in the unstructured tail of the substrate. **b)** Example trace for the substrate processing assay monitoring SspB-mediated degradation of non-ubiquitinated I27$^{V15P}$ substrate. Two binding events are shown, with the first one (left) being non-productive and the second (right) leading to degradation. **c)** Representative trace for the conformational change assay during I27$^{V13/15P}$ substrate degradation by SspB-fused ΔRpn10 proteasome. The substrate-processing dwells (blue shading) for this substrate are characterized by their lack of transitions to the low-FRET s1 state. **d)** Representative trace for the conformational change assay during $I27^{V13/15P}$ substrate degradation by the SspB-fused Rpn5-mutant proteasome shows a substrate-processing dwell that also lacks s1 excursions. **e)** Representative trace for the conformational change assay during ubiquitin-dependent degradation of the $I27^{V15P}$ substrate degradation. The substrate-processing dwell (red shading) shows brief s1 excursions during the unfolding phase, but excursions are absent during translocation (hatched; see also Fig. 2e, Supp. Fig. 9, 12).

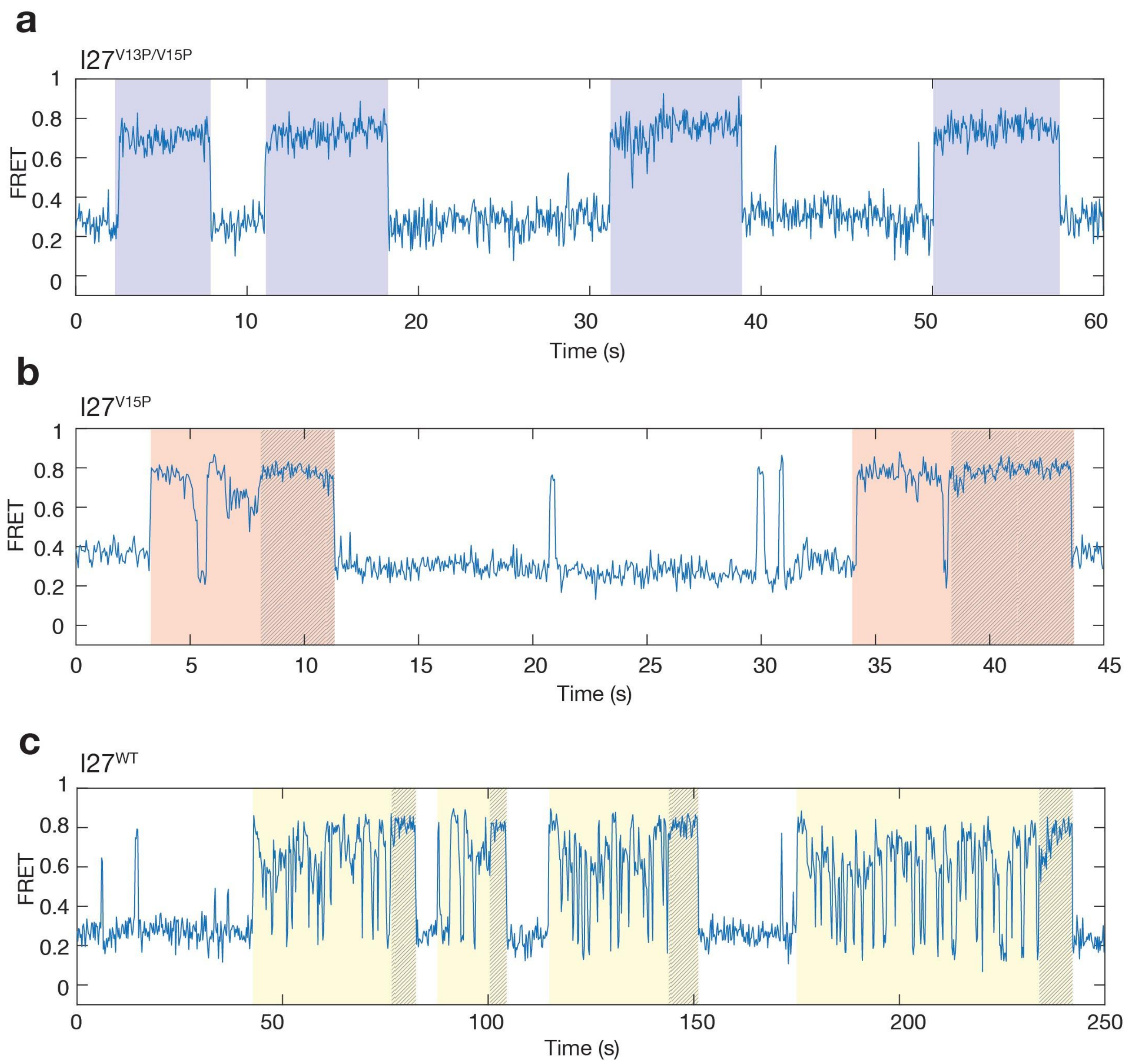


**Supplementary Figure 9: Substrate processing dwells and s1-state excursions. a)** Representative trace for the conformational change assay with four consecutive substrate processing dwells (blue shading) for SspB-mediated degradation of I27$^{V13/15P}$ substrate. Dwells for this thermodynamically labile substrate do not exhibit pronounced transitions to the low-FRET s1 state. **b)** Representative trace with two consecutive substrate-processing dwells (red shading) for SspB-mediated degradation of I27$^{V15P}$ substrate. Dwells for this substrate show s1 transitions early, potentially during unsuccessful unfolding attempts, but lack them during the last 3-5 s (hatched), which likely represents translocation of the unfolded polypeptide. **c)** Representative trace with four consecutive substrate-processing dwells (yellow shading) for SspB-mediated degradation of wild-type I27 substrate. Dwells for this thermodynamically most stable substrate exhibit frequent excursions to the low-FRET s1 state, except for the last 3-5 s (hatched).

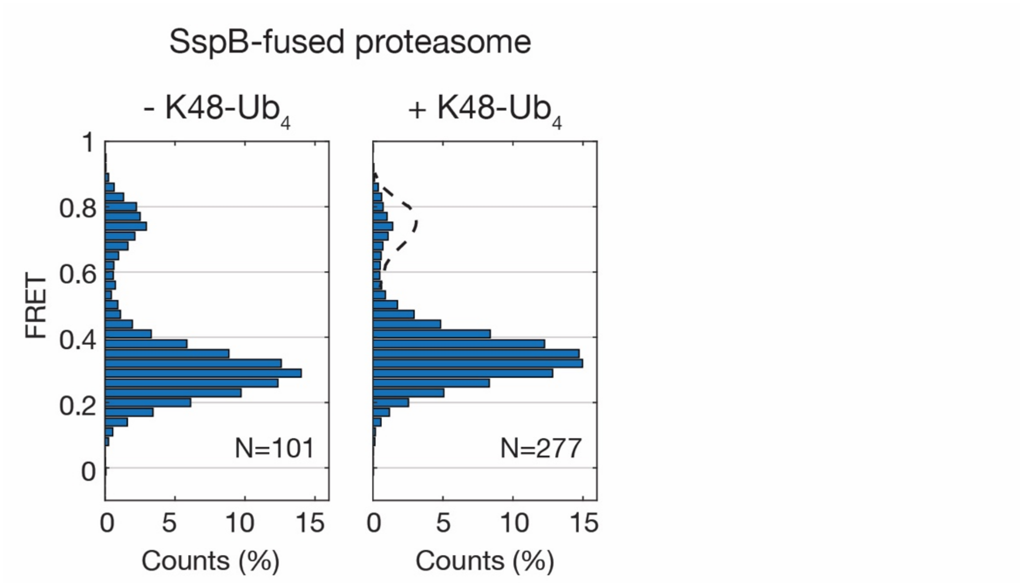


**Supplementary Figure 10: Conformational dynamics of the SspB-fused proteasome are modulated by ubiquitin chains similar to wild-type proteasome.** FRET-state histograms for SspB-fused proteasomes in the absence (left) and presence (right) of K48-linked tetra-ubiquitin chains.

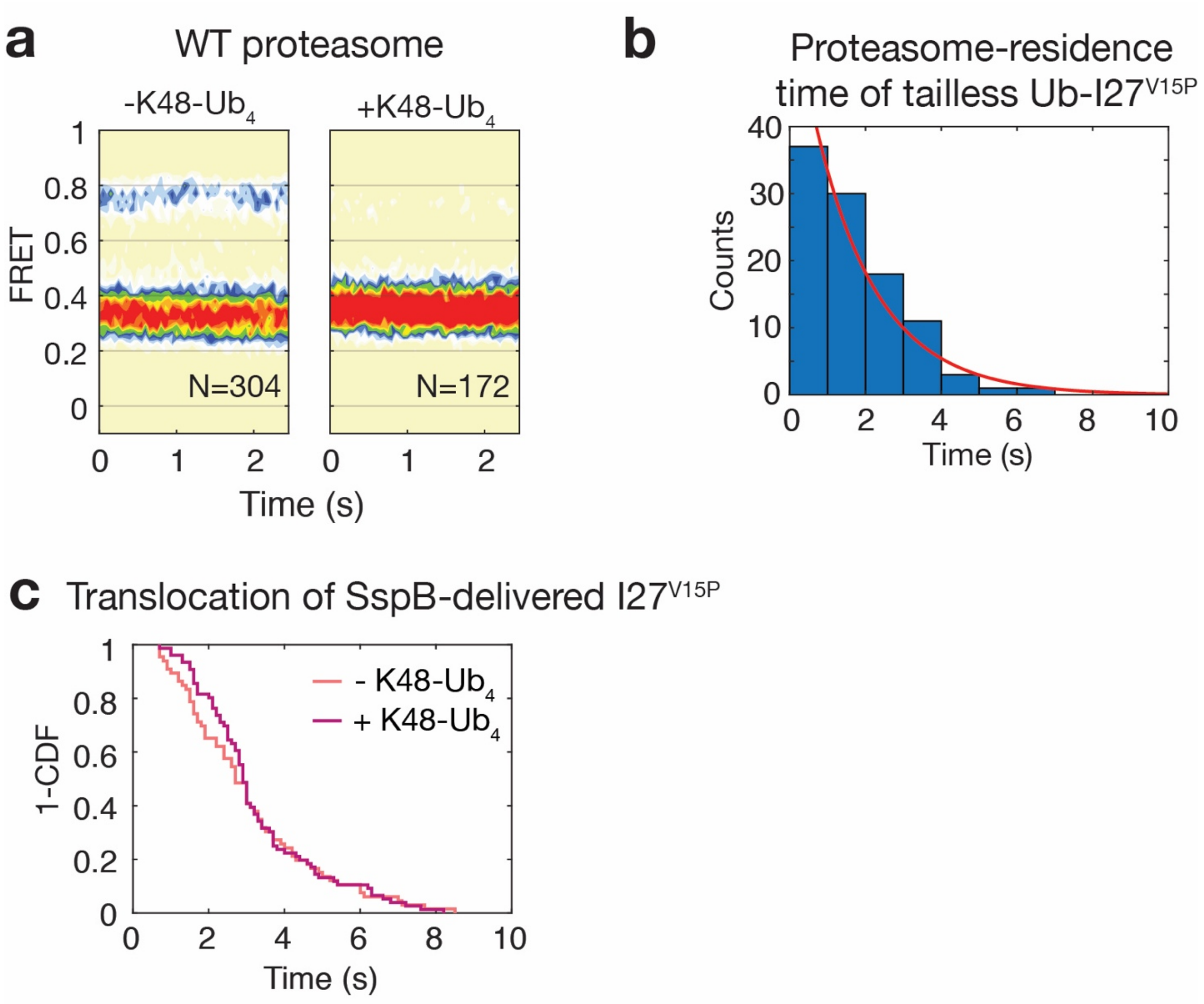


**Supplementary Figure 11: Proteasome interactions with ubiquitin. a)** Contour plots for the FRET-state occupancies of wild-type proteasome in the absence (left) and presence (right) of K48-Ub4. This is an alternative representation of the data shown in figure 3b. **b)** Dwell-time histogram for the residence time of a ubiquitinated tailless $I27^{V15P}$ substrate on the proteasome (N = 101). This substrate is unable to engage with the ATPase motor and therefore only exhibits non-productive binding, indicated by short events with intermediate FRET value. The histogram was fit to a single exponential (red line) to determine the time constant $\tau_{off}$ = 0.61 ± 0.12 s for the dissociation of an ubiquitinated substrate from proteasomal receptors (error is 95% confidence interval of the fit). **c)** FRET-decay times in the substrate-processing assay for the degradation of $I27^{V15P}$ show no difference in the absence versus presence of K48-Ub4 (N = 66 and 76 respectively).

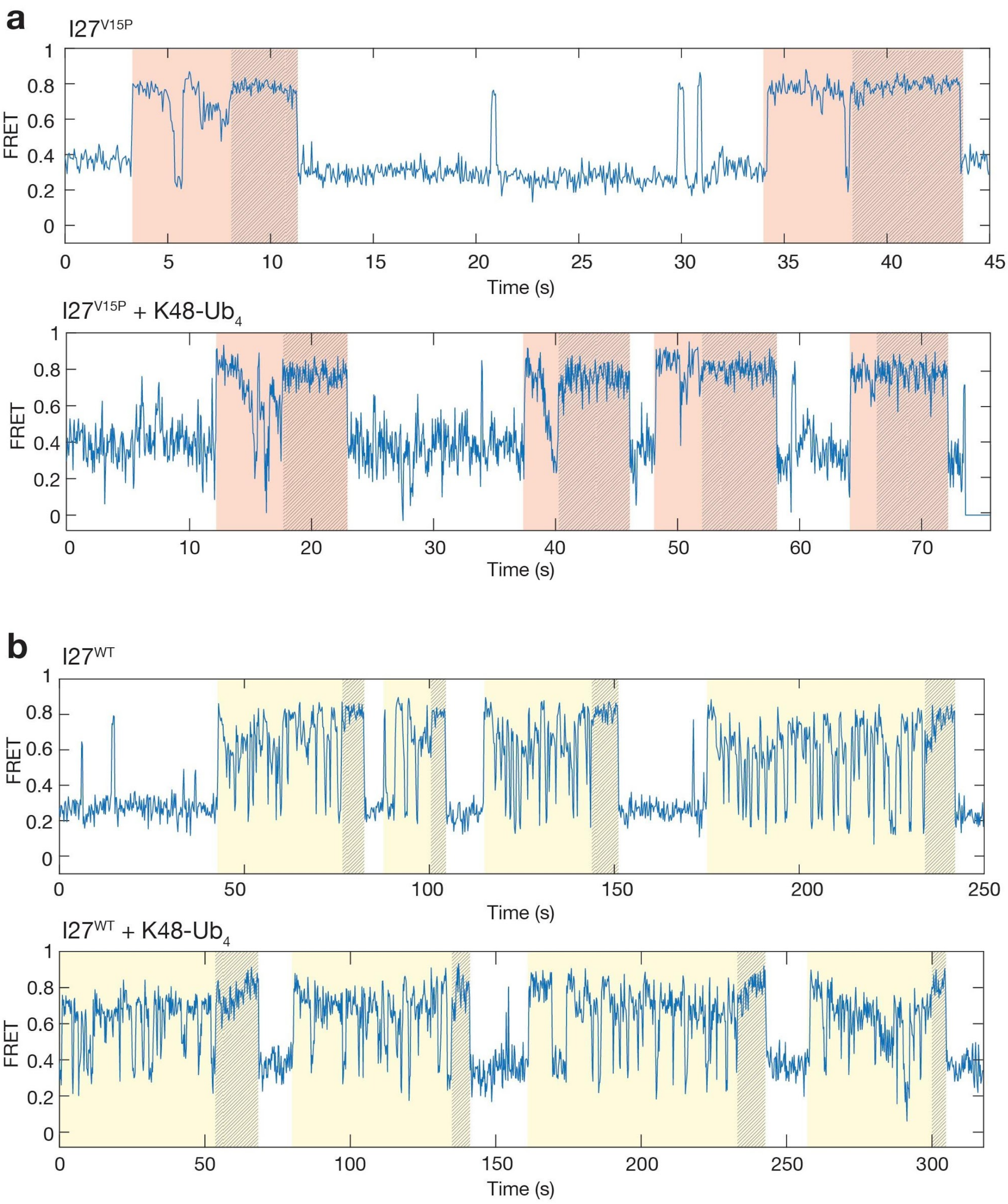


**Supplementary Figure 12: No obvious modulation of proteasome conformational dynamics by ubiquitin during substrate degradation. a)** Representative traces for the conformational change assay monitoring SspB-mediated I27$^{V15P}$ degradation in the absence (top) and presence (bottom) of K48-Ub$_4$ show no significant difference in the frequency or duration of s1 excursions during substrate-processing dwells. **b)** Similarly, processing dwells during the SspB-mediated I27$^{WT}$ degradation exhibit no discernable differences in conformational switching behavior between the absence (top) and presence (bottom) of K48-Ub$_4$.

| Substrate | Mean □ s.e.m. (s) | N |
|---|---|---|
| Ub-I27$^{V15P}$ + WT proteasome | 1.8 □ 0.1 | 80 |
| Ub-I27$^{V15P}$ + Rpn11$^{V80A}$ proteasome | 1.7 □ 0.1 | 74 |
| I27$^{V15P}$ + SspB-fused proteasome | 3.4 □ 0.3 | 77 |
| I27$^{V15P}$ + SspB-fused proteasome + K48-Ub$_4$ | 2.2 □ 0.2 | 102 |

**Supplementary Table 1: Substrate tail-insertion and engagement kinetics.** Listed are the mean values of the time constants for tail insertion and engagement of either ubiquitinated or SspB-delivered I27$^{V15P}$ substrate, in the absence and presence of ubiquitin chains, as determined by the FRET-based tail insertion and processing assay. N values indicate the number of analyzed events.

| Proteasome variant | $k_{s1}$ ($s^{-1}$) | $k_{non\text{-}s1}$ ($s^{-1}$) |
|---|---|---|
| WT proteasome | 1.2 □ 0.1 | 4.5 □ 0.1 |
| WT proteasome + K48-Ub$_4$ | 0.4 □ 0.1 | 4.2 □ 0.1 |
| Rpn10-ΔUIM proteasome | 0.9 □ 0.1 | 2.1 □ 0.1 |
| SspB-fused proteasome | 0.8 □ 0.1 | 4.2 □ 0.1 |
| SspB-fused proteasome + K48-Ub$_4$ | 0.4 □ 0.1 | 3.8 □ 0.1 |
| Rpn5-mutant proteasome | 2.5 □ 0.1 | 4.2 □ 0.1 |

**Supplementary Table 2: Rates of proteasome conformational switching.** Rates for the conformational transitions from the s1 to non-s1 states ($k_{s1}$) and from non-s1 to s1 states ($k_{non\text{-}s1}$) were determined by Hidden-Markov-modeling of the results from the FRET-based conformational change assay for wild-type and various mutant proteasomes in the absence and presence of unanchored ubiquitin chains.

| Substrate | Mean □ s.e.m. (s) | N |
|---|---|---|
| I27$^{V13P/V15P}$ | 7.3 □ 0.2 | 102 |

| | | |
|---|---|---|
| I27$^{V13P/V15P}$ + K48-$Ub_4$ | 7.0 □ 0.1 | 111 |
| I27$^{V15P}$ | 10.7 □ 0.6 | 85 |
| I27$^{V15P}$ + K48-$Ub_4$ | 9.0 □ 0.3 | 113 |
| I27$^{WT}$ | 47 □ 4 | 67 |
| I27$^{WT}$ + K48-$Ub_4$ | 40 □ 4 | 66 |

**Supplementary Table 3: Substrate processing dwells.** Given are the mean values for the duration of high-FRET processing dwells measured by the FRET-based conformational change assay during the SspB-mediated degradation of I27-substrate variants in the absence and presence of unanchored ubiquitin chains. The N values indicate the number of analyzed events.
